# Characterization of Vlf1 as a regulator of lipophagy

**DOI:** 10.64898/2026.08.11.744108

**Authors:** Zacharias Fakih, Claudia Cavarischia-Rega, Brian Russell Glück, Simeon Reichert, Parijat Dutta, Olga Beresh, Maya Schuldiner, Boris Maček, Doron Rapaport, Kai Stefan Dimmer

## Abstract

Lipid droplets (LDs) are unique organelles, surrounded by a phospholipid monolayer. They are present in most eukaryotic cells including the unicellular model organism *S. cerevisiae*. LDs store neutral lipids which serve as precursors for amphipathic membrane lipids and as an energy reserve. Loss of LDs in *S. cerevisiae* results in multiple cellular defects impairing lipid homeostasis and the biogenesis and function of other organelles. Here, we find that the expression levels of many proteins in isolated mitochondrial fractions are altered in cells that cannot synthesize neutral lipids and therefore lack LDs. In addition, among several downregulated proteins, we identified the previously uncharacterized Ylr001c (which we name Vlf1 for *V*acuolar *L*ipophagy *F*actor *1*). We show that Vlf1 is glycosylated and, in contrast to some previous reports, is actually localized to the vacuole. Furthermore, we demonstrate that changes in Vlf1 expression alter growth sensitivity to rapamycin, and detected a physical interaction of Vlf1 with Atg15, a lipase involved in autophagy. Additionally, we observe higher levels of autophagy/lipophagy in the absence of Vlf1 and a reduction upon overexpression of the protein. Taken together, the effects on lipohagy by Vlf1 makes it, according to our knowledge, the first vacuolar lipophagy regulator identified in *S. cerevisiae*.

## Introduction

Lipid droplets (LDs) are key organelles for lipid storage and metabolic regulation in eukaryotic cells. By storing neutral lipids such as triacylglycerols (TAG) and sterol esters (SE), LDs buffer fluctuations in nutrient availability and help protect cells from lipotoxic stress (Bohnert, 2026; Henne, 2023; Mathiowetz and Olzmann, 2024). Beyond their storage function, LDs engage in extensive functional and physical interactions with other organelles, including the endoplasmic reticulum, mitochondria, and the vacuole/lysosome (Herker et al., 2021; Schuldiner and Bohnert, 2017). Consequently, loss of LDs has profound effects on cellular physiology, impacting membrane homeostasis, stress resistance, and energy metabolism (Czabany et al., 2007; Graef, 2018; Henne and Cohen, 2026).

In *Saccharomyces cerevisiae* (from here on yeast), loss of LD formation results in pleiotropic phenotypes, reflecting the broad role of LDs in cellular organization and metabolism. Previous studies have shown that LD-deficient cells display altered lipid composition (Sorger et al., 2004), increased sensitivity to stress (Petschnigg et al., 2009), and defects in metabolic adaptation (Garbarino et al., 2009). Given the central role of mitochondria in energy conversion, it has been proposed that mitochondria may be particularly affected by LD loss. Indeed, changes in mitochondrial morphology and distribution were reported (Bischof et al., 2017).

The molecules stored within LDs are either utilized for synthesis of other membrane lipids or for energy conversion. Two major pathways are responsible for the degradation/mobilization of LDs. In lipolysis, LDs stay intact and their cargo is degraded by lipases, which are intrinsic LD proteins. During autophagy, on the other hand, whole LDs are degraded. Autophagy is a fundamental cellular quality-control mechanism that helps maintain homeostasis under stress conditions (Kotani and Nakatogawa, 2026). When organelles become damaged or dysfunctional, cells can activate selective forms of autophagy, to remove and recycle these defective compartments (Onishi et al., 2021; Wang et al., 2026). This adaptive response not only restores metabolic balance but also supports cell survival in conditions where normal organelle function is compromised (Picca et al., 2023). A special form of autophagy, the so called lipophagy mobilizes stored lipids for energy production and helps maintain cellular lipid homeostasis, particularly under nutrient stress (Fairman and Ouimet, 2022; Kounakis et al., 2019). Lipophagy in yeast is a special form of microautophagy where LDs are directly engulfed by the vacuole, yet there are also reports that LD proteins rely on the common pathway of macroautophagy (from here on autophagy) via autophagosome formation (Schott et al., 2022; van Zutphen et al., 2014).

In yeast, mitochondria and peroxisomes cooperate as key hubs of lipid utilization, with β-oxidation occurring in peroxisomes and the resulting metabolites being used by mitochondria for energy conversion (Hiltunen et al., 2003; Pascual-Ahuir et al., 2017; Shai et al., 2018). In addition, mitochondria form contact sites with LDs and other organelles, facilitating lipid exchange and metabolic communication (Bohnert, 2020; Dimmer and Rapaport, 2017; Enkler and Spang, 2024). These observations led us to hypothesize that loss of LDs would trigger adaptive changes in the mitochondrial proteome of yeast, potentially revealing factors involved in lipid homeostasis or compensatory metabolic pathways. To test this hypothesis, we set out to identify mitochondrial proteins whose abundance is altered in cells lacking LDs.

Among the putative mitochondrial proteins affected by LD depletion, we identified Ylr001c which had been previously reported in systematic studies to have a mitochondrial and/or vacuolar localization. We renamed the protein Vlf1 (for *V*acuolar *L*ipophagy *F*actor *1*) and show that it is actually a vacuolar protein and physically interacts with Atg15, the vacuolar phospholipase B (Watanabe et al., 2023). We additionally demonstrate that Vlf1 exerts regulatory influence on autophagic processes, most prominently on the autophagy of lipid droplets (lipophagy).

## Results

### Yeast lacking lipid droplets have reduced levels of the uncharacterized protein Ylr001c

It was previously observed that the loss of lipid droplets (LDs) leads to changes in mitochondrial morphology (Bischof et al., 2017). Therefore, we assayed the proteome of mitochondria to uncover proteins that are differentially expressed in yeast cells lacking LDs. We performed a SILAC based mass spectrometry on crude mitochondrial fractions from either control cells or cells lacking the four enzymes required for triacylglycerol (TAG) and sterolester (SE) synthesis, and therefore entirely lacking lipid droplets (ΔLD). More than 1430 proteins were identified for both cell types, including 671 mitochondrial proteins. From all the proteins with altered expression levels shown in the volcano plot of Figure 1A (see complete data set in Supplementary Table S1), we selected those that were identified as mitochondrial (Supplementary Table S2 provides an overview of such proteins whose expression levels changed by at least 1.5-fold). Most candidates were localized to the inter membrane space (IMS) or the mitochondrial matrix. Since we were interested in factors that potentially contribute to the interaction between LDs and mitochondria, we focused on the uncharacterized protein Ylr001c. Considering our observations described below, we renamed the protein Vlf1 (for *V*acuolar *L*ipopagy *F*actor *1*).

**Fig. 1.**
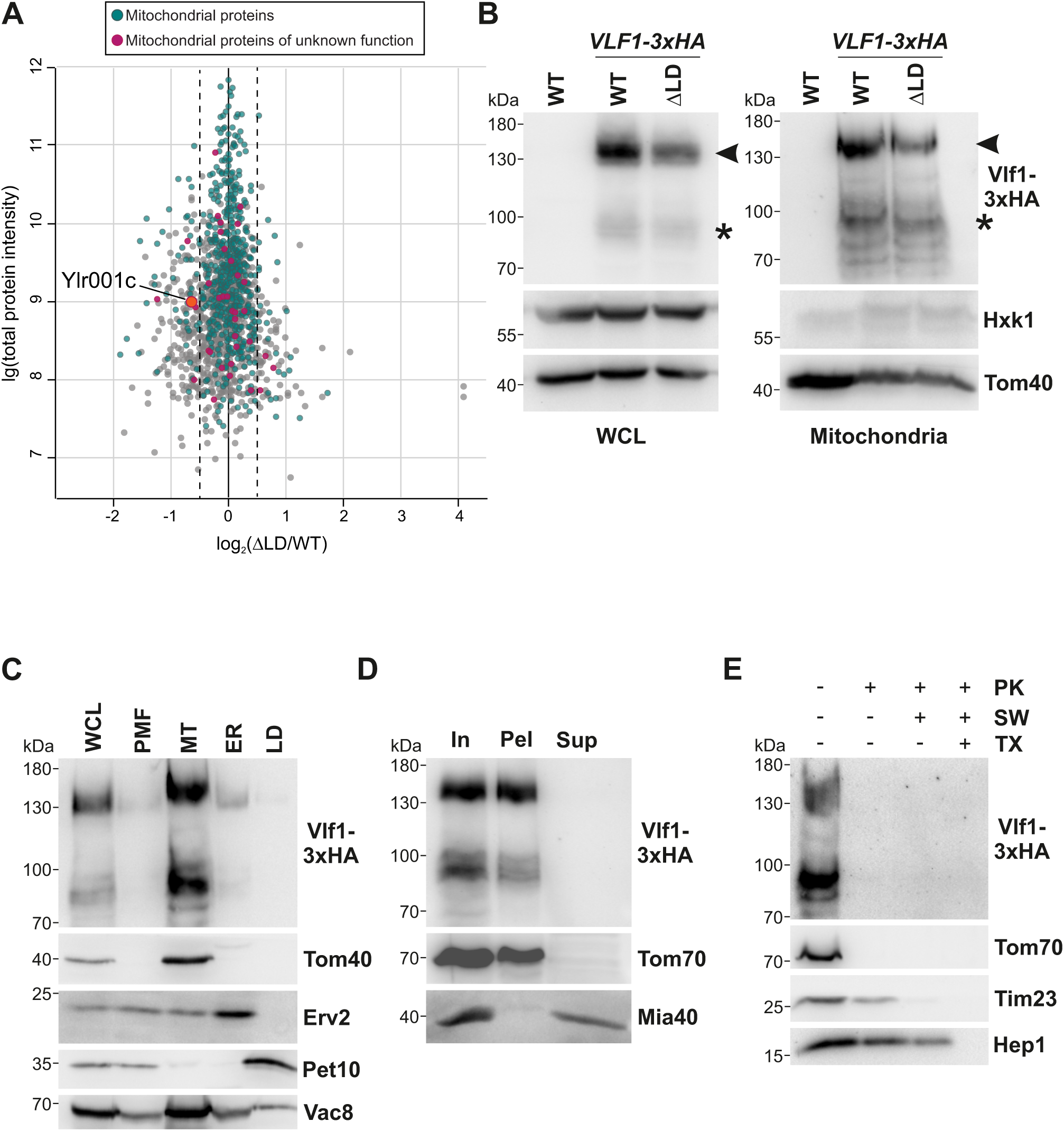
Vlf1 is a membrane protein downregulated in cells lacking lipid droplets. **(A)** Volcano plot showing expression changes of proteins between control (WT) and ΔLD cells. Annotated mitochondrial proteins and mitochondrial proteins of unknown function are shown by the indicated colors. Ylr001c (Vlf1) as putative mitochondrial protein of unknown function is highlighted separately. **(B)** Vlf1 levels are reduced in ΔLD cells. Whole cell lysates (WCL) or crude mitochondrial fractions were obtained from either control (WT) or ΔLD cells harboring endogenously HA-tagged Vlf1. The samples were analyzed by western blotting with antibodies against the HA-tag, as well as Tom40 (mitochondrial marker) and Hxk1 (cytosolic marker). The arrowhead marks the dominant band of approximately 130 kDa, the asterisk the 90 kDa form. **(C)** Vlf1 is present in the crude mitochondrial fraction in subcellular fractionation analysis. Equal amounts of protein of the indicated fractions (PMF, post mitochondrial fraction; MT, mitochondrial fraction; ER, microsomal fraction; LD, lipid droplet fraction) were analyzed by western blotting with the indicated antibodies. Erv2 (ER marker); Pet10 (LD marker); Vac8 (vacuole marker). **(D)** Vlf1 is an integral membrane protein. Crude mitochondria isolated from Vlf1-3xHA expressing cells (In, Input) were subjected to alkaline extraction. The pellet (Pel) and supernatant (Sup) fractions were analyzed by western blotting with antibodies against the indicated proteins. Tom70, an integral MOM protein exposed to the cytosol; Mia40, a soluble IMS protein. **(E)** Vlf1 faces the cytosol. Crude mitochondria as in (D) were treated with proteinase K (PK) under different conditions. Mitochondria were kept intact, the MOM was ruptured by hypo-osmolar swelling (SW), or mitochondria were lysed by the addition of Triton X-100 (TX). Samples were analyzed by western blotting with the indicated antibodies. Tim23, a mitochondrial inner membrane protein exposed to the IMS; Hep1 (matrix marker).

To verify that Vlf1 levels are reduced in ΔLD cells, we obtained by tetrad dissection a ΔLD yeast strain that contains C-terminally HA-tagged Vlf1. Indeed, whole-cell lysates (WCL) from these yeast cells showed reduced Vlf1-3xHA levels in comparison to those in control cells, and crude mitochondrial fractions of the same cells showed similar reduction in the protein amounts (Fig. 1B). Of note, we detected two specific bands for Vlf1-3xHA, a dominant one of ca. 130 kDa (Fig. 1B, arrowhead) and a less pronounced one of approx. 90 kDa (Fig. 1B, asterisk). As in our study, Vlf1 was previously found in mitochondrial isolations in several systematic studies (Reinders et al., 2006; Schulte et al., 2023; Sickmann et al., 2003). On the other hand, several previous high-throughput microscopic approaches (Huh et al., 2003; Yofe et al., 2016) report a predominantly vacuolar localization of the protein. Furthermore, the sequence homolog of Vlf1 from *S. pombe,* Fsc1, was shown to be localized to the vacuole (Sun et al., 2013). In agreement with non-mitochondrial localization, Fsc1 also shows sequence and structural homology with human fasciclin proteins involved in extracellular matrix structure, cell-cell adhesion, paracrine signaling, and endocytosis (Seifert, 2018). In support of a vacuolar localization also of Vlf1, AlphaFold2 suggests that in addition to a putative transmembrane domain near its C-terminus and four Fas1 domains, the protein contains several putative N-glycosylation sites that are not found in mitochondrial proteins (Suppl. Fig. S1) (Jumper et al., 2021). To resolve the discrepancy regarding Vlf1’s localization, we initially performed subcellular fractionation of yeast cells by differential centrifugation and found that Vlf1, like the mitochondrial protein Tom40, is enriched in the mitochondrial fraction. However, the vacuolar marker protein Vac8 was also present in this fraction, precluding precise subcellular localization at this stage (Fig. 1C). Vlf1 contains a putative transmembrane domain close to its C-terminus (Suppl. Fig. S1). Hence, we performed a carbonate extraction experiment to test whether it is a membrane-embedded protein. Indeed, Vlf1 is found in the pellet fraction, like other integral membrane proteins, e.g. the mitochondrial outer membrane (MOM) protein Tom70 and in contrast to the soluble IMS protein Mia40 (Fig. 1D). To determine the orientation of Vlf1 in the membrane, we treated crude mitochondria with proteinase K under different conditions. Upon proteolytic digestion, the signal for Vlf1-3xHA was already lost in intact mitochondria and therefore was also absent when either the MOM was ruptured by osmotic swelling (SW) or all membranes were solubilized by the addition of the detergent Triton X-100 (TX)(Fig. 1E). Considering the prediction of a single transmembrane domain, this result supports the notion that C and N-termini of the protein face different compartments of the cell, probably with the C-terminus facing the cytosol.

### Vlf1 is a vacuolar protein

As mentioned above, we primarily observed upon immunodecoration a 130-kDa species instead of the expected ∼100-kDa band based on the predicted molecular mass of the protein tagged with three HA epitopes. Since previous systematic proteome analysis identified 13 different N-glycosylation sites for Vlf1 (https://glycosmos.org/glycoproteins/Q07895) (Zielinska et al., 2012) we investigated whether this difference is caused by glycosylation of the protein. To that goal, we treated whole-cell lysates of yeast harboring the tagged protein with the glycosidases PNGase or EndoH. Indeed, upon this treatment, Vlf1 behaved similarly to the known N-glycosylated protein Carboxypeptidase 1 (Prc1), and we found a reduction of the molecular mass of Vlf1 to a species of ca. 100 kDa (Fig. 2A).

**Fig. 2.**
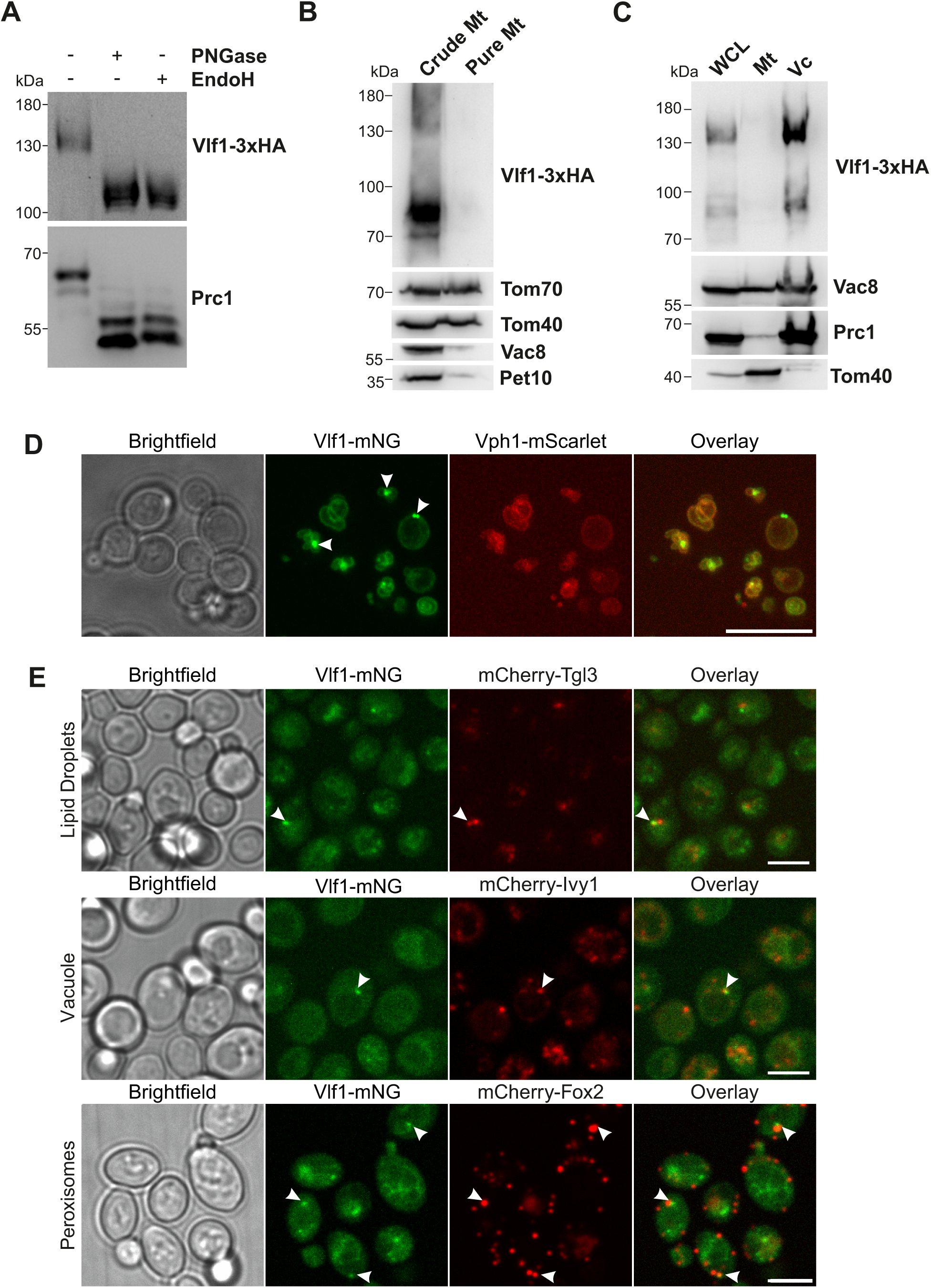
Vlf1 is a vacuolar protein forming punctate structures. **(A)** Vlf1 is glycosylated. Whole cell lysates of cells harboring HA-tagged Vlf1 were left untreated or treated with the glycosidases PNGase F or Endo H. Samples were analyzed by western blotting with antibodies against the HA-tag and Prc1 (a glycosylated vacuolar protein). **(B)** Vlf1 is absent in highly purified mitochondrial fractions. Mitochondrial membranes isolated by either spheroplasting and differential centrifugation (Crude Mt) or further purified by sucrose gradient centrifugation (Pure Mt) were analyzed by western blotting with the indicated antibodies. **(C)** Vlf1 is found in a vacuolar membrane fraction. Vacuolar membranes and mitochondria were isolated from lysed yeast cells harboring Vlf1-3xHA. Equal protein amounts of lysed cells (WCL), isolated mitochondria (Mt) or vacuolar (Vc) membranes were analyzed by western blotting with the indicated antibodies. **(D)** Fluorescently labelled Vlf1 is located to the vacuole. Yeast cells expressing endogenously tagged Vlf1-mNG and Vph1-mScarlet (a vacuolar marker) were analyzed by fluorescence microscopy. Punctate structures formed by Vlf1 in a subset of cells are marked by white arrowheads. Scale bar: 10 µm. **(E)** Vlf1-mNG punctae colocalize with several punctae-forming proteins of different organelles. Heterozygous diploid yeast cells expressing endogenously tagged Vlf1-mNG and the depicted mCherry-labelled protein were analyzed by fluorescence microscopy. Colocalizing punctate structures are marked by white arrowheads. Scale bar: 5 µm.

Since glycosylation is characteristic of proteins that follow the secretory pathway rather than mitochondrial proteins, this observation prompted us to reevaluate the subcellular localization of Vlf1. Hence, we asked whether Vlf1 would be detected also in highly pure mitochondria obtained by sucrose density step gradient centrifugation. This additional purification step resulted in almost complete loss of the Vlf1-3xHA signal similarly to the behaviour of known non-mitochondrial proteins like Vac8 (vacuole) and Pet 10 (LDs) (Fig. 2B). We therefore applied a previously published protocol for vacuole purification by subcellular fractionation using a Ficoll density gradient (Kagohashi et al., 2023). Figure 2C shows that Vlf1-3xHA indeed behaves like the vacuolar marker proteins Vac8 and Prc1 and in contrast to the mitochondrial marker Tom40.

To further confirm the vacuolar localization, we genomically tagged Vlf1 with a green fluorescent tag (mNeonGreen; Vlf1-mNG) and analyzed the cells by fluorescence microscopy. We found that the protein co-localizes with the mScarlet-tagged vacuolar protein Vph1 (Fig. 2D). Of note, employing N-terminally GFP-tagged Vlf1, it was previously suggested that the N-terminus of the protein faces the lumen of the vacuole (Baruch et al., 2025). Together with the result of our proteolytic analyses (Fig. 1E), these observations imply that Vlf1 is a vacuolar transmembrane protein with its N-terminus facing the lumen of the vacuole and its C-terminus facing the cytosol. Interestingly, in some cells Vlf1-mNG showed additional punctate structures colocalizing with the vacuolar membrane (Fig. 2D, arrowhead).

To better characterize the nature of Vlf1 punctate structures, we employed a systematic high content screening approach to identify proteins that colocalize with Vlf1. For this purpose, a library of approximately 1,300 yeast strains, each expressing a distinct mCherry-tagged protein with previously reported punctate distribution in the cell, was crossed with a strain expressing Vlf1-mNG. The resulting heterozygous diploids, co-expressing Vlf1-mNG and the respective candidate protein, were analyzed by fluorescence microscopy. Since the *S. pombe* homologue Fsc1 formed similar punctate structures under starvation conditions, the doubly labelled cells were cultivated under glucose and nitrogen starvation conditions. Among the screened strains, 22 exhibited clear partial colocalization with Vlf1-mNG (Supplementary Table S3). These candidates predominantly included vacuolar, peroxisomal, and LD-associated proteins (each comprising 23% of the total number of hits), along with a smaller fraction of mitochondrial proteins (13%). Figure 2E illustrates examples of colocalizations with proteins residing in the three most prominent organelles mentioned above. The observed overlaps may reflect a functional link between vacuolar Vlf1 and other organelles.

### Vlf1 interacts with Atg15 and forms an Atg15-dependent protein complex

Next, we wanted to identify Vlf1’s interaction partners and analyze if it is a subunit of a higher molecular weight complex. To address this question, we first verified that Vlf1-3xHA can be efficiently immunoprecipitated using an antibody specific for the HA-epitope (Fig. 3A). Next, the elution fraction enriched in Vlf1-3xHA was analyzed by mass spectrometry for co-precipitating proteins. Although we observed 516 proteins in control samples unspecifically bound we found overall more proteins to be pulled down (579 proteins) in Vlf1-3xHA immunoprecipitates. Of note, the same proteins were found with high reproducibility between all three biological replicates (Suppl. Fig. S2A, B). Consistent with successful enrichment, Vlf1 displayed increased intensity, peptide counts, and sequence coverage in the immunoprecipitated samples compared with controls (Suppl. Fig. S2C). Several proteins were significantly enriched in the bound fraction, the most prominent protein identified was Atg15 (Fig. 3B), a phospholipase B required for autophagy (Kagohashi et al., 2023; Watanabe et al., 2023). Atg15 was also strongly enriched with respect to intensity, peptide counts, and sequence coverage in the Vlf1-3xHA samples compared with controls (Suppl. Fig. S2D). Of note, this interaction was also reported in a high throughput mass spectrometry-based approach aimed at analyzing the complete interactome of *S. cerevisiae* proteins (Michaelis et al., 2023).

**Fig. 3.**
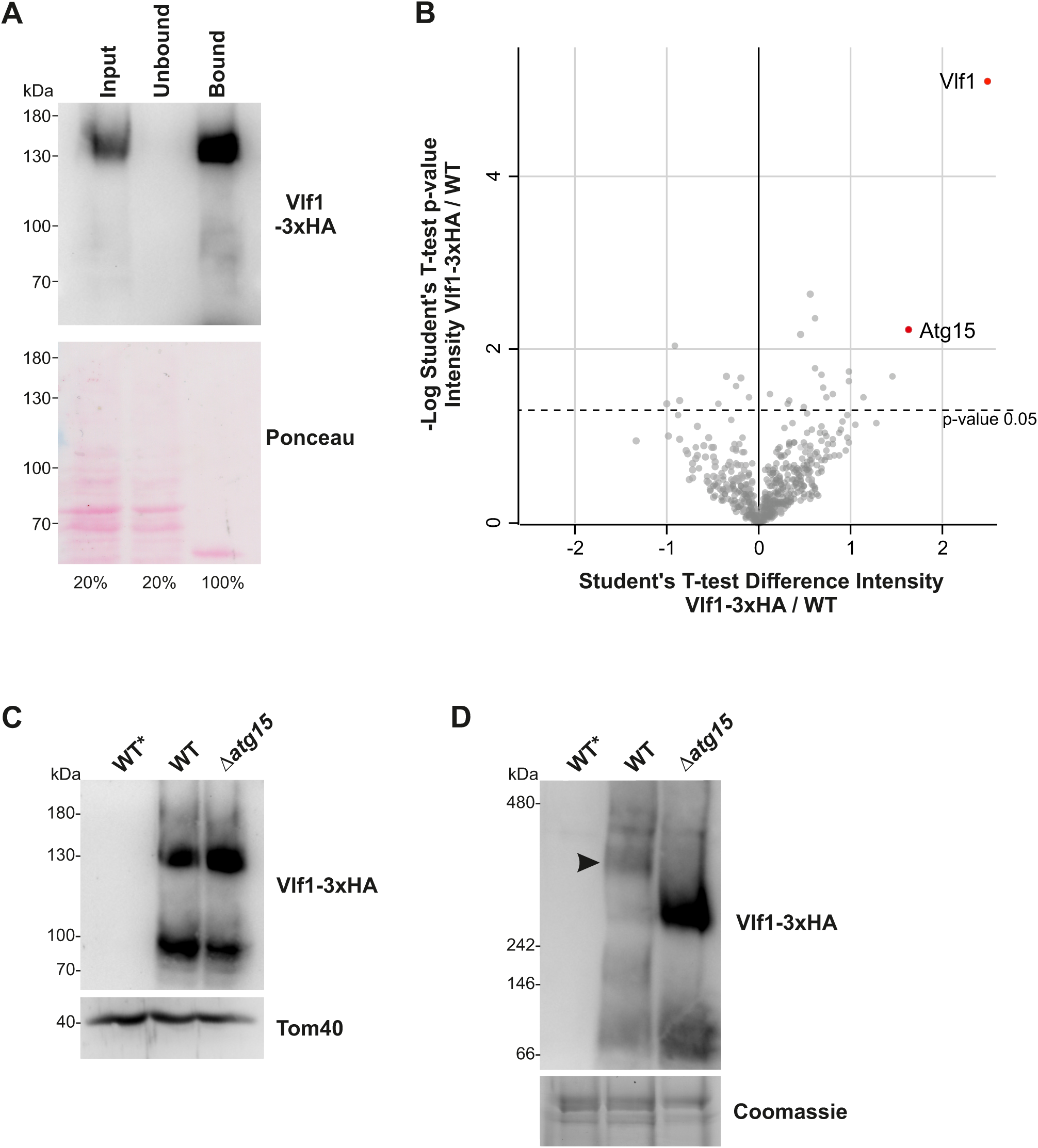
Vlf1 physically interacts with the vacuolar phospholipase B Atg15. **(A)** Immunoprecipitation of Vlf1. Vlf1-3xHA was pulled down from whole cell lysates of yeast cells. Input (20%) and unbound proteins (20%) as well as bound proteins (100%) were analyzed by western blotting with an anti-HA antibody. The lower panel shows the Ponceau staining of the membrane. **(B)** Vlf1-3xHA co-precipitates with Atg15. Volcano plot of changes in intensity for co-precipitated proteins between control (WT) and Vlf1-3xHA harboring cells. Vlf1 and Atg15 are highlighted in orange and red, respectively. **(C)** Steady-state levels of Vlf1-3xHA do not change upon loss of Atg15. Equal amounts of crude mitochondria isolated from either control (WT) or *atg15*Δ cells containing Vlf1-3xHA were analyzed by western blotting with antibodies against the HA-tag and Tom40 as loading control. **(D)** Vlf1 forms higher molecular weight complexes dependent on Atg15. Crude mitochondria containing Vlf1-3xHA were solubilized with TritonX-100 (Tx-100) and subjected to BN-PAGE followed by western blotting with an antibody against the HA-tag. A part of the Coomassie stained membrane is shown as loading control. The arrowhead marks a higher molecular weight complex of approximately 350 kDa.

Since interacting proteins often affect each other’s stability, we next analyzed the steady state levels of Vlf1 in the *ATG15* deletion background. However, we found that the expression levels of Vlf1-3xHA were unaltered compared to control cells suggesting that Atg15 is not affecting the stability of Vlf1(Fig. 3C). Next, to determine whether Vlf1 is a component of a higher molecular weight complex that might also contain Atg15, we analyzed crude mitochondrial fractions containing Vlf1-3HA by BN-PAGE (for unknown reasons vacuolar fractions could not be analyzed by BN approaches). Vlf1 was detected in control samples in several bands but mainly as part of a higher molecular weight complex of approximately 350 kDa (Fig. 3D, arrowhead). The detection of Vlf1 in several bands is in agreement with its broad migration pattern as detected in a recent high throughput complexome study (Schulte et al., 2023). Supporting an Atg15-Vlf1 physical interaction, we found that the migration of Vlf1 changed in the absence of Atg15, and the tagged protein appeared in a smaller complex of approximately 250 kDa.

To analyze whether the two factors genetically interact, we constructed a double deletion mutant *atg15*Δ/*vlf1*Δ which did not show any growth defect on different carbon sources (Suppl. Fig. S3A). Of note, the deletion of *VLF1* in cells lacking LDs moderately enhanced the growth retardation of these cells at elevated temperature (37°C) (Suppl. Fig. S3A). In contrast, we found that overexpression of Vlf1 had no impact on the growth of control, *atg15*Δ or ΔLD cells under all tested conditions (Suppl. Fig. S3B). In summary, our results suggest that the autophagy factor Atg15 and Vlf1 interact with each other and are part of a protein complex at the vacuolar membrane.

### Vlf1 puncta formation is impacted by autophagy conditions

Importantly, a functional connection of Vlf1 with the autophagy factor Atg15 is further strengthened by the fact that the punctate structures of Vlf1 are absent under any condition in the *ATG15* deletion mutant (Fig. 4A, second and fourth row). Next, we asked whether the sub-cellular distribution of Vlf1 would be affected by autophagy promoting conditions. Indeed, adding the mTOR inhibitor rapamycin that induces autophagy to logarithmically growing cells led to a change in the distribution of Vlf1. The number of cells that contained punctate structures, as well as the number of structures per cell dramatically increased (Fig. 4A, row 1 compared to row 3). Of note, a similar behavior was observed for the structural homologue from *S. pombe*, Fsc1 (Sun et al., 2013). These findings strongly suggest a role for Vlf1 in autophagic processes.

**Fig. 4.**
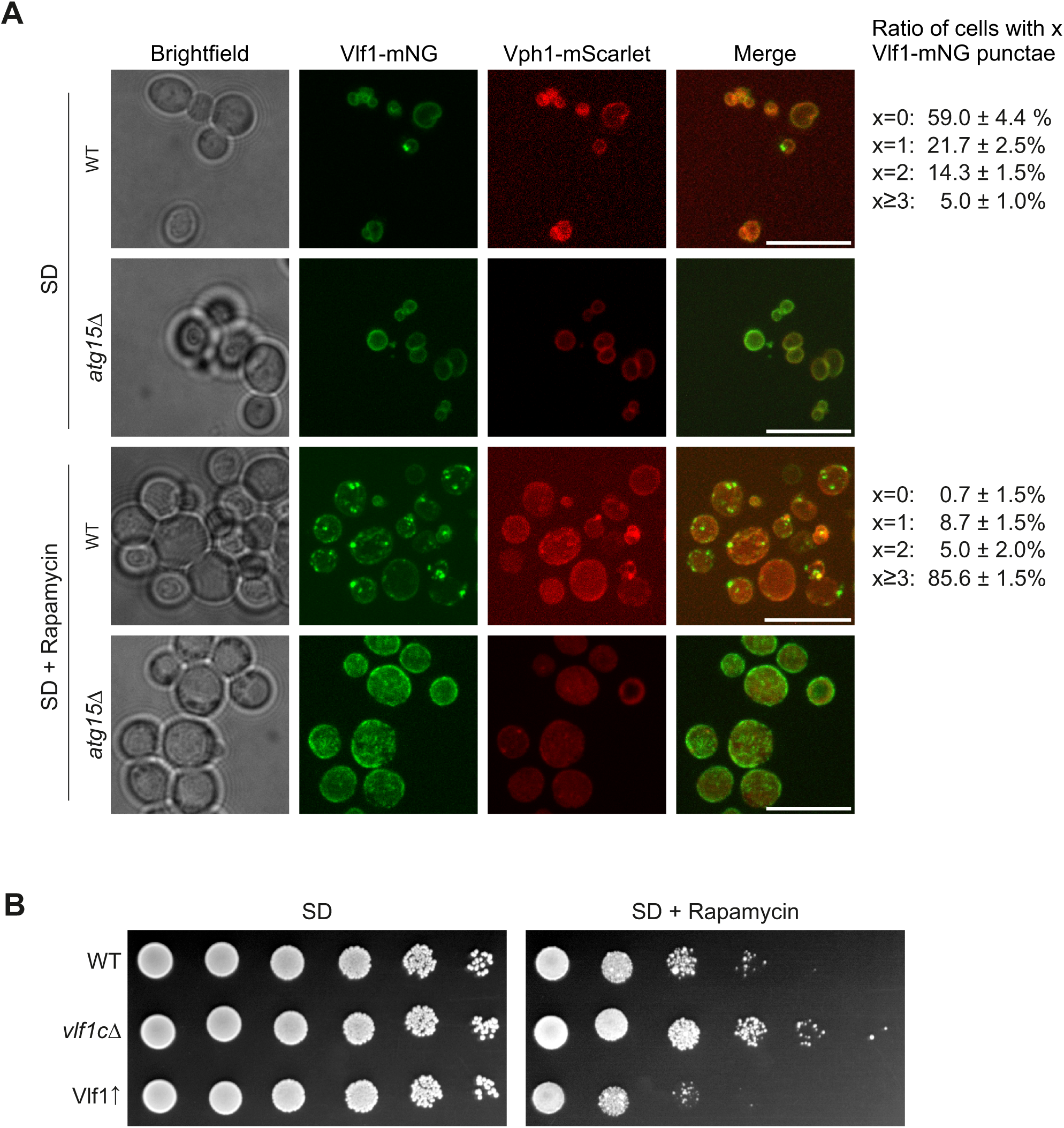
Vlf1 influences the sensitivity of yeast cells to rapamycin. **(A)** The punctate structures containing Vlf1 depend on Atg15 and are affected by rapamycin. Control (WT) and *atg15*Δ cells expressing endogenously labelled Vlf1-mNG and Vph1-mScarlet were grown to mid-logarithmic phase in the presence or absence of rapamycin and analyzed by confocal fluorescence microscopy. Representative images are shown. Scale bar: 10 µm. A quantitative analysis of n=3 independent experiments each comprising at least 100 cells of each yeast strain is provided on the right side of the image series. **(B)** Vlf1 levels influence the growth of yeast cells in the presence of rapamycin. Control (WT), *vlf1*Δ and Vlf1 overexpressing (Vlf1↑) cells were grown to logarithmic phase and spotted on either SD or SD + rapamycin (2.5 ng/mL) plates in a 1:5 dilution series. Plates were incubated at 30°C.

### Vlf1 levels impact autophagy

Next, we asked whether levels of Vlf1 influence growth when yeast cells are treated with rapamycin. Notably, whereas under normal growth conditions neither the deletion nor the overexpression of Vlf1 affect growth, adding rapamycin to the growth medium changed this behavior. In the presence of rapamycin, cells lacking Vlf1 grew significantly better than control cells and in contrast cells overexpressing Vlf1 from a strong synthetic promoter (*P_7tet1_*, (Azizoglu et al., 2021)) showed a slight, yet reproducible growth defect (Fig. 4B).

We then asked whether Vlf1 might be involved in general autophagy, or in specific forms like mitophagy and/or lipophagy. To this end, we first analyzed whether global autophagy was compromised in cells lacking or overexpressing Vlf1. We examined the behavior of the fusion protein GFP-Atg8 in *vfl1*Δ and Vlf1 overexpressing cells, compared to control cells, under different stimuli known to induce autophagy. We quantified the extent of autophagy by the standard approach of tracking the formation of cleaved, free GFP species. We observed elevated free GFP levels, especially in stationary phase and upon glucose withdrawal, in *vlf1*Δ cells, and a reduction when Vlf1 was overexpressed. Rapamycin addition and nitrogen starvation resulted only in minor changes, yet with the same tendencies, in cells with altered Vlf1 levels compared to control cells (Fig. 5A). As control for the autophagy-related appearance of free GFP, this species was not detected in logarithmic grown cells or upon the deletion of the autophagy promoter, Atg1 (Fig. 5A). Taken together, these results indicate that under certain conditions, namely stationary phase and glucose depletion, the loss of Vlf1 facilitates autophagy, whereas overexpression has an inhibitory effect under certain conditions, namely stationary phase and glucose depletion.

**Fig. 5.**
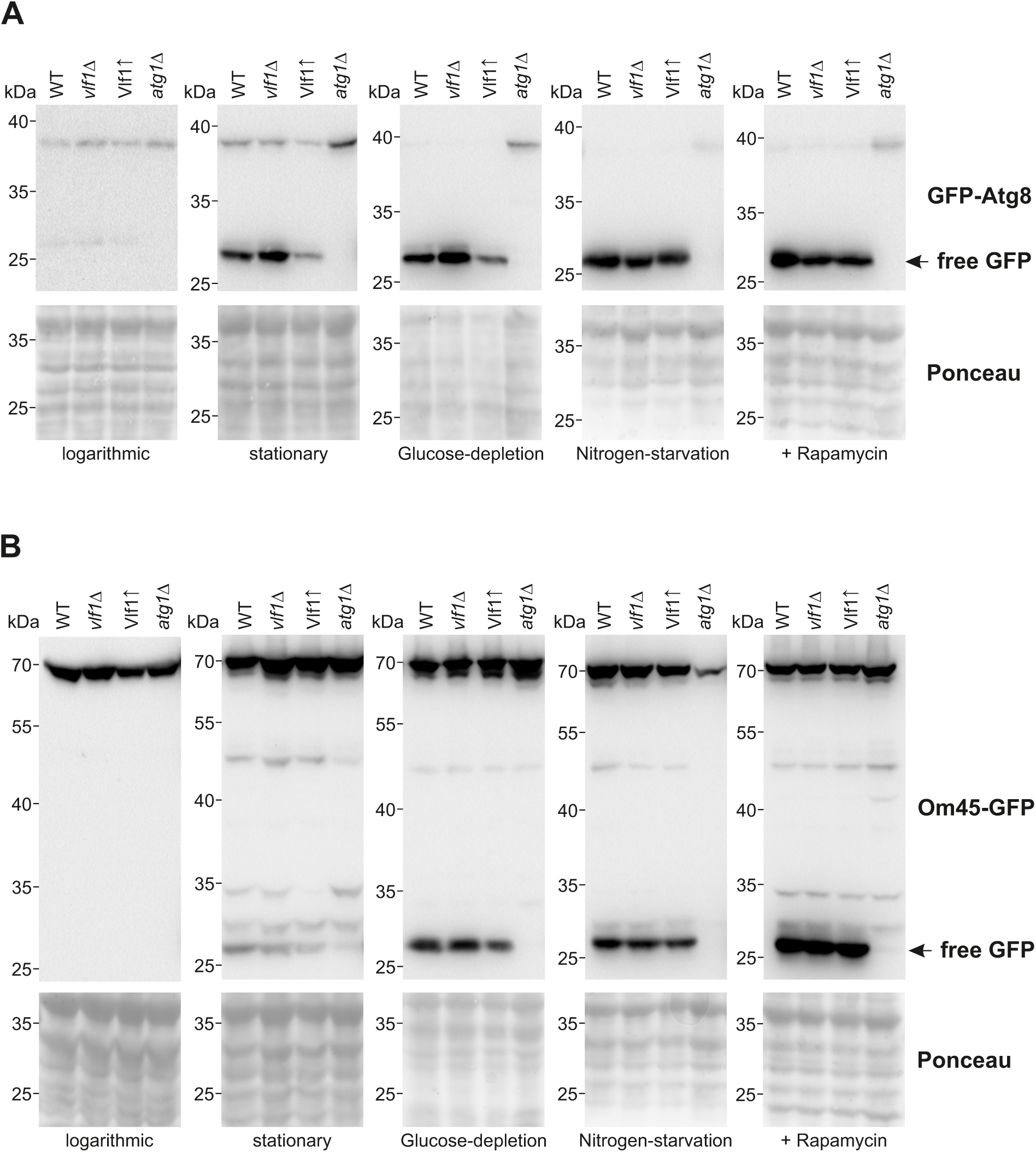
Vlf1 levels modulate certain types of autophagy. **(A)** Control (WT), *vlf1*Δ, Vlf1↑ or *atg1*Δ cells expressing plasmid-borne GFP-Atg8 were cultured in SD medium and were treated in one of the following ways: (i) grown to mid logarithmic growth phase (logarithmic), (ii) grown overnight to stationary phase (stationary), (iii) shifted for 16 h to S medium with 0.001% (w/v) glucose (Glucose depletion), (iv) shifted for 16 h to SD-N medium (Nitrogen starvation) or (v) grown for 16 h on SD medium containing 100 ng/mL rapamycin (+Rapamycin). Afterwards, whole cell lysates were analyzed by western blotting with an antibody against GFP. The migration of free GFP is indicated. The Ponceau staining of the Western Blot is shown as loading control. A representative experiment of n≥3 repetitions is shown. **(B)** The same strains as in (A) expressing endogenously GFP labelled OM45 were treated as described in (A). Whole cell lysates were prepared and analyzed by western blotting as described in (A).

To analyze mitophagy, we then analyzed free GFP levels derived from the MOM protein OM45-GFP upon autophagy induction by different stimuli. The levels of free GFP were not influenced by elimination or high levels of Vlf1 under all inducing conditions (Fig. 5B), suggesting that Vlf1 is likely not involved in mitophagy regulation.

### Vlf1 levels affect lipid droplet number and lipophagy

Both glucose depletion and stationary phase are known conditions to induce also lipophagy. Therefore, we wondered whether loss of Vlf1 influences the quantity of LDs in yeast cells. Indeed, loss of Vlf1 led to a decrease in number of LDs compared to corresponding control cells during logarithmic growth, as detected by fluorescence microscopy with the LD marker Pet10-mNG (Fig. 6A, left panel). Quantitative flow cytometry analysis of the intensity of fluorescence signal of Pet10-mNG per cell shows a statistically significant decrease in mean cellular fluorescence in the absence of Vlf1 (Fig. 6A, right panel). This difference is also present in stationary-phase cells (Fig. 6B). Additionally, most lipid droplets in stationary *vlf1*Δ cells appear to surround the vacuole whereas control cells have LDs spread out in the cytosol (Fig. 6B).

**Fig. 6.**
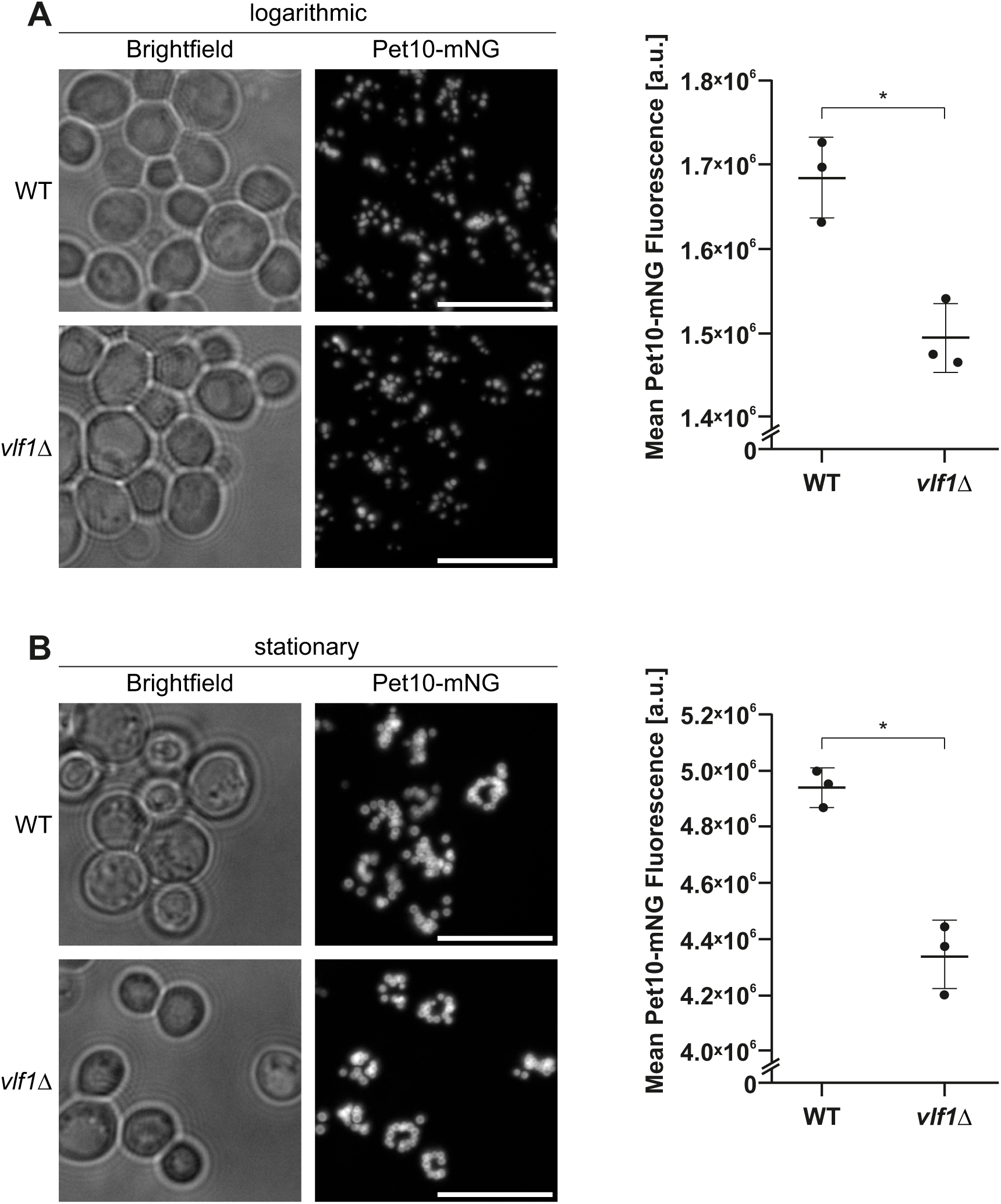
Vlf1 levels influence lipid droplet numbers per cell. **(A)** Control (WT) and *vlf1*Δ cells expressing endogenously mNG-labelled Pet10 (Pet-mNG) were grown to mid-logarithmic phase. Left panel: representative fluorescence microscopy images (scale bar = 10 µm). Right panel: the cells described above were analyzed by flow cytometry. The mean fluorescence per cell was measured and the average with standard deviation of three biological replicates (n=3) is shown. *, *P*<0.05 (unpaired t-test, two-tailed) **(B)** Same as in (A) for stationary cells.

To study lipophagy in detail, we analyzed its extent under the same autophagy-inducing conditions introduced in the previous section, by monitoring the formation of free GFP from the LD protein Faa4 fused to GFP (Fig. 7A). The amount of free GFP present in these cells reflects the overall degree of lipophagy. Under all conditions, loss of Vlf1 led to a higher amount of free GFP, whereas increased Vlf1 levels showed the opposite effect. To further strengthen these findings, we also monitored this process in stationary cells over longer periods of time and obtained the same trend (Fig 7B).

**Fig. 7.**
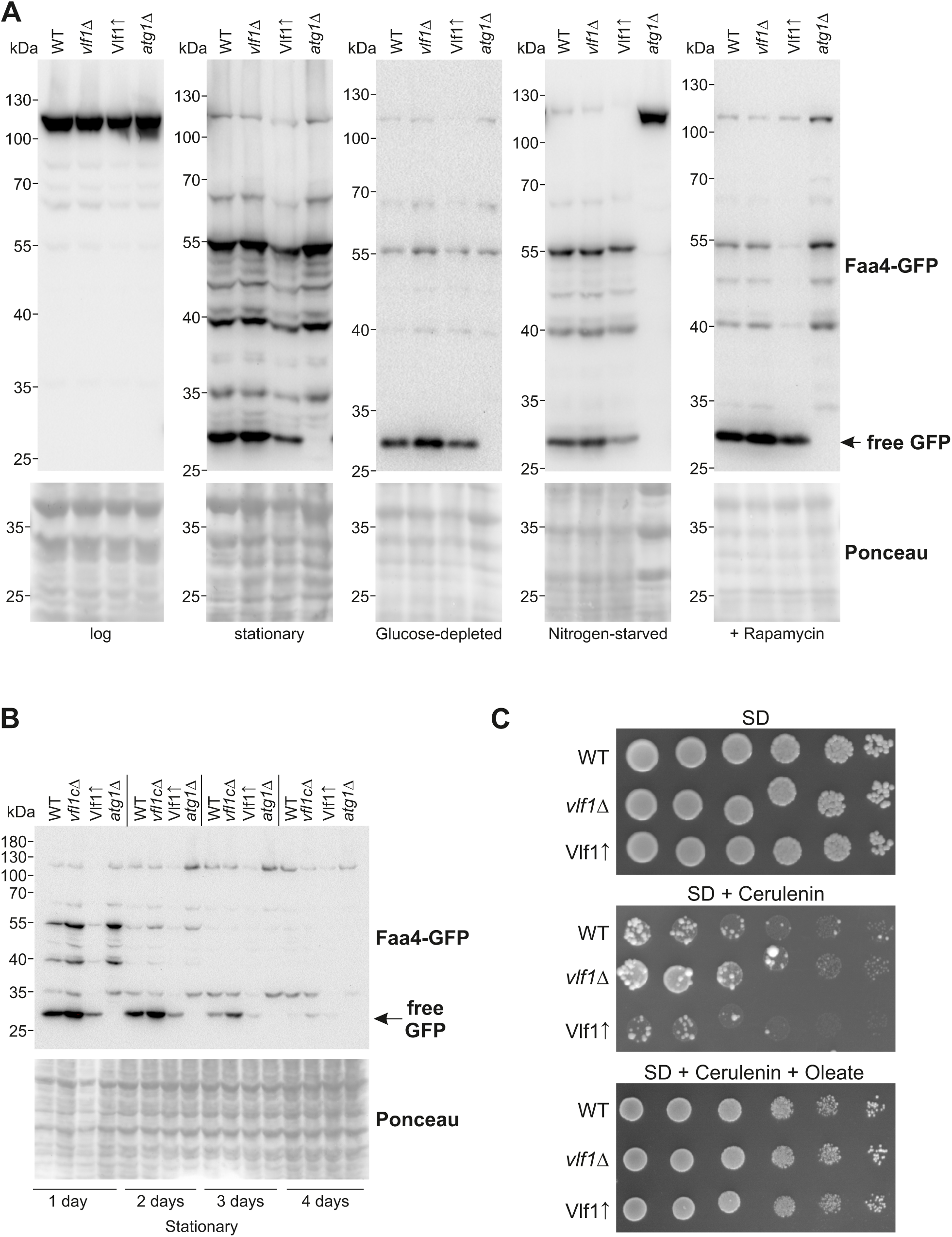
Vlf1 levels influence lipophagy levels under different conditions. **(A)** The same cells as in Figure 5A endogenously expressing Faa4-GFP were cultured in SD medium to the mid logarithmic growth phase and then further treated as described in the legend to Fig. 5A. **(B)** The same strains as in (A) were kept for the indicated number of days in stationary phase and whole cell lysates were analyzed as described in (A). **(C)** The designated cells were grown to logarithmic phase and spotted in a 1:5 dilution series on plates containing either SD, SD + 0.25 µg/mL cerulenin or SD + 0.25 µg/mL cerulenin + oleate. Plates were incubated at 30°C.

We conclude that Vlf1 appears to inhibit lipophagy. Consequently, one would expect that Vlf1 interferes with the mobilization of LD components, such as fatty acids stored as triacylglycerides. Supporting this assumption, we found that growth of yeast cells upon addition of cerulenin, an inhibitor of *de novo* fatty acids synthesis, is enhanced in the absence of Vlf1, whereas cells with higher levels of Vlf1 show reduced growth when fatty acid synthesis is inhibited (Fig. 7C, middle panel). This observation suggests that the loss of fatty acid synthesis caused by cerulenin can be partially compensated by increased mobilization of stored fatty acids via enhanced lipophagy in cells lacking Vlf1. Of note, we observed the appearance of several rapidly growing colonies for all three strains (Fig. 7C, middle panel) consistent with a strong selection for endogenous suppressors previously reported to be induced by cerulenin treatment (Inokoshi et al., 1994). As expected, supplementation of the medium with the fatty acid oleate in addition to cerulenin eliminated the need for *de novo* fatty acid synthesis and rescued the growth phenotype of all tested strains (Fig. 7C, lower panel). Taken together, our findings strongly support a regulatory inhibitory function for Vlf1 in lipophagy.

## Discussion

Lipid droplets are the main storage compartment for neutral lipids in eukaryotic cells. These lipids can be mobilized by enzymatic lipolysis or lipid droplet autophagy (lipophagy). In yeast, in contrast to higher eukaryotes, lipophagy can also occur via microautophagy (through direct invagination of the vacuole membrane). Here, we describe a novel factor for autophagy, particularly for lipophagy that interacts with Atg15, the only autophagy-related phospholipase known so far. Atg15 was primarily implicated in the degradation of autophagic bodies after autopahgosome-vacuole fusion (Watanabe et al., 2023).

We identified Vlf1 as a protein with reduced levels in yeast cells lacking LDs. Although we initially found the protein in crude mitochondrial fractions, our additional biochemical and microscopic analysis showed that it is a vacuolar membrane protein. These apparently contradicting observations reflect disagreement in previous reports. On the one hand, Vlf1 was found in several systematic studies to be enriched in crude mitochondrial fractions (Reinders et al., 2006; Schulte et al., 2023; Sickmann et al., 2003), on the other hand fluorescence microscopy suggested that the protein is located in the vacuole (Huh et al., 2003; Yofe et al., 2016). The fact that Vlf1 was not detected in a more recent proteomic study employing highly pure mitochondria provides further support for its vacuolar localization (Vogtle et al., 2017). We show that Vlf1 is part of a higher-molecular-weight complex. In addition, Vlf1 can be detected in punctate structures by fluorescence microscopy. Both the punctate structures and the specific oligomeric form strongly rely on the presence of its interaction partner, Atg15. We speculate that the formation of these structures in the vacuolar membrane could explain Vlf1’s presence in crude mitochondrial membrane fractions. Assuming that Vlf1 complexes reside in higher-density vacuolar membrane patches, vesicles containing these patches could form when the vacuole is ruptured during subcellular fractionation. These vesicles could co-sediment with mitochondria due to their similar density. Another potential explanation for the presence of Vlf1 in crude mitochondrial fractions is the possibility that a subpopulation of the protein is involved in contact sites between mitochondria and vacuole.

Collectively, we present four pieces of evidence for the vacuolar localization of Vlf1: (i) Vlf1 cannot be detected anymore in pure mitochondrial fractions after purification by density gradient centrifugation. (ii) Upon using a protocol dedicated to isolating purified vacuoles, Vlf1 behaves like the vacuolar protein Vac8. (iii) We show that Vlf1 is N-glycosylated, a posttranslational modification that occurs in the ER and Golgi and (iv) our microscopic analyses reveal a clear vacuolar distribution of mNG-tagged Vlf1, with the protein predominantly staining the vacuolar membrane and forming a few punctate structures.

These punctate structures likely represent Vlf1-enriched vacuolar membrane subdomains. They might be interpreted as the fluorescent counterpart of the Vlf1-containing complexes detected by BN-PAGE. Interestingly, these punctae increase in number if the autophagy inducer rapamycin is added and depend on the presence of the autophagy lipase Atg15.

Vlf1’s localization is indicative for a function at the vacuole, the site of autophagy, and mirrors the vacuolar membrane localization of the *S. pombe* structural homolog Fsc1 (Sun et al., 2013). Fsc1 localizes to the vacuole membrane and forms starvation-induced punctae, which are partially dependent on Atg1, Atg11, and Atg13 (Sun et al., 2013). The fact that Vlf1 also forms punctate structures, and that these depend on Atg15, suggests that puncta formation of Vlf1-like proteins may be a conserved, autophagy-related feature that could mark sites of membrane remodeling during organelle turnover. However, although these punctate structures are dependent on the presence of Atg15, which was shown to have a single N-terminal transmembrane domain (Marquardt et al., 2023), further experiments are necessary to determine their molecular composition and their physiological function.

Strong support to the role of Vlf1 in regulating lipophagy comes from the observation that, cells lacking Vlf1 on the other hand are less sensitive to rapamycin, whereas cells with higher Vlf1 levels are more sensitive. Since rapamycin induces autophagy, the reduced sensitivity of Vlf1-deficient cells could be explained by the removal of an inhibitory brake on autophagy/lipophagy. In the absence of Vlf1, autophagy induction by rapamycin can take place more efficiently, allowing cells to better cope with rapamycin-induced stress. Conversely, higher Vlf1 levels may restrain autophagy/lipophagy, making cells more vulnerable to rapamycin. This interpretation is consistent with our observations showing that Vlf1 acts as an inhibitory regulator of autophagy/lipophagy under glucose withdrawal and during stationary phase. In these conditions, loss of Vlf1 leads to increased lipophagic flux, as evidenced by elevated free GFP levels in GFP-Atg8 and Faa4-GFP assays, concomitant with a reduction in LD number. Overexpression of Vlf1 has the opposite effect, reducing free GFP generation. The cerulenin phenotype further supports this inhibitory role. Cerulenin inhibits fatty acid synthase, preventing de novo fatty acid synthesis and forcing cells to rely on lipid stores and lipophagy for fatty acid supply. Cells lacking Vlf1 are less sensitive to cerulenin, consistent with increased lipophagic flux that allows more efficient mobilization of fatty acids from LDs. Conversely, cells with higher Vlf1 levels are more sensitive to cerulenin, likely because Vlf1-mediated inhibition of lipophagy limits fatty acid release from LDs when synthesis is blocked. Thus, Vlf1 appears to function as an inhibitory regulator of lipophagy, restraining lipid droplet turnover under glucose withdrawal and stationary phase, and its removal facilitates autophagy/lipophagy induction by rapamycin and improves fatty acid supply.

Although Vlf1 and the *S. pombe* protein Fsc1 are structural homologs and both localize to the vacuolar membrane with puncta formation under autophagy inducing conditions, their proposed functions appear to differ. Fsc1 has been reported to be crucial for autophagosome–vacuole fusion, acting as a positive mediator of this step in *S. pombe* (Sun et al., 2013). In contrast, our data suggests that Vlf1 functions as an inhibitory regulator. This divergence may reflect functional specialization of Vlf1-like proteins across fungi, where a conserved vacuolar membrane scaffold or complex has been adapted to either promote or inhibit distinct steps in autophagy/lipophagy depending on the organism and cellular context.

Overall, we introduce Vlf1 as a novel vacuolar regulator that specifically inhibits lipophagy while leaving selective mitophagy unaffected. Although its precise molecular mechanism remains to be fully deciphered, our data position Vlf1 as a modulator of LD turnover. Future work will need to address whether Vlf1 also influences other selective autophagy pathways, such as pexophagy, Golgi-phagy, ER-phagy, or nucleophagy, and whether it acts as a dedicated lipophagy brake or as part of a broader organellophagy regulatory network.

## Material and Methods

### Yeast cell culture

Yeast strains were grown in standard rich medium (YP) or synthetic medium (S) with either glucose, galactose, glycerol, ethanol or oleate as the carbon source.

Transformation of yeast cells was achieved using the lithium acetate method. For cells transformed with plasmids, synthetic media with appropriate selection marker(s) were used. For drop dilution assays, yeast cells were cultured to an OD_600_ of 1.0 and serially diluted in five-fold increments. Aliquots (5 µL) of the serially diluted cultures were spotted onto the corresponding solid medium, and the cells were incubated at either 30 or 37°C.

To induce autophagy by nitrogen starvation, mid-logarithmic cells were harvested, washed three times in S-N medium (S-medium with the required carbon source without ammonium sulfate and without amino acids) and resuspended in S-N medium. To induce lipophagy by glucose depletion, mid-logarithmic cells were harvested, washed three times with S-D medium (S-medium containing the required amino acids and 0.001 % (w/v) glucose), and resuspended in S-D medium.

### Yeast genetic manipulation

Most deletion and genomically tagged strains were made by homologous recombination. Gene-specific primers were used to amplify deletion cassettes from the pFA6a vector series (Hentges et al., 2005; Janke et al., 2004) unless described otherwise. The correct integration of the deletion cassette was confirmed by PCR using primers specific to the genes of interest.

Yeast strains harboring mNeonGreen tagged Pet10 were generated by CRISPR/Cas9-based techniques. To this end, a plasmid carrying Cas9 under the estradiol-inducible promoter and a region in which the sgRNAs can be integrated into was used (pZF041) so that the sgRNA can be transcribed simultaneously with Cas9. The sgRNAs were designed with the sequence 5’-GCG CCG GCT GGG CAA CAC CTT CGG GTG GCG AAT GGG ACC NNNNNNNNNNNNNNNNNNNN GTT TTA GAG CTA GAA ATA GCA AGT TAA AAT AAG GC-3’ where the N_20_ sequence was replaced by the target sequence. To define the targeting sequence, a PAM sequence corresponding to NGG (where N could be any base) was chosen in proximity to the region to be manipulated. The first 20 bp directly upstream of the PAM sequence were taken as the targeting sequence. The so designed sgRNA and its reverse complement were ordered as DNA oligonucleotides from Eurofins Genomics. Both oligonucleotides were mixed 1:1 and heated for 5 min at 95 °C. After cooling and double strand DNA formation of the oligonucleotides, linearized pZF041 (*Bsa*I and *Afl*II) was added, and the mixture transformed into yeast cells. Next, single transformed colonies were streaked in parallel on YPD+G418 and YPD+G418+β-estradiol plates. Cells from colonies growing significantly slower on YPD+G418+β-estradiol were transformed with the template DNA for homologous recombination. The template for the insertion of the tag contained the desired sequence and in addition 5’ and 3’ overhangs of 45 bp homologous to the insertion sites and were generated by PCR or annealing of ordered oligonucleotides. For the transformation in the Cas9 carrying cells, cells were grown in YPD+G418 to mid logarithmic phase, β-estradiol (10 µM final conc.) was added, and cells were incubated for additional 3 h, followed by transformation with the template for homologous recombination. Clones carrying the desired genetic modification were selected for by plating and growth on YPD+G418+β-estradiol plates.

Yeast strains used in this study are described in Supplementary Table S4.

### Recombinant DNA techniques

The ORF of *YLR001C* with all *Eco*RI sites eliminated, was purchased from Eurofins Genomics as plasmid pEX-A258-YLR001C. To express HA-tagged protein, the ORF without the Stop-codon was subcloned with the restriction enzymes *Eco*RI and *Xma*I into the yeast expression vector pYX142 to generate pYX142_YLR001C-HA.

Primers and plasmids used in this study are listed in Supplementary Tables S5 and S6, respectively.

### Isolation of mitochondria

Mitochondria were isolated from yeast cells using a previously published protocol based on differential centrifugation (Daum et al., 1982). Yeast cultures (2-5 L) were grown at 30°C and harvested at an OD_600_ of 0.8 by centrifugation (3000xg, 5 min, RT). The cell pellets were washed, weighed and resuspended in resuspension buffer (100 mM Tris, 10 mM DTT). The cell suspension was incubated at 30°C for 10 min after which the cells were harvested as above. The cell pellets were washed with a spheroplasting buffer (1.2 M sorbitol, 20 mM potassium phosphate, pH 7.2). Next, the cell pellets were resuspended in zymolyase-containing spheroplasting buffer (5 mg zymolase per g cell pellet) and incubated at 30°C for at least 1 h with constant shaking. The spheroplasts were harvested (2000xg, 5 min, 2°C), resuspended in homogenization buffer (0.6 M Sorbitol, 1 mM EDTA, 1 mM PMSF, 0.2% (w/v) fatty-acid free BSA, 10 mM Tris, pH 7.4), and dounced 15 times. The lysed spheroplasts were first subjected to two clarifying steps (2000xg, 5 min, 2°C) and then to a high-speed centrifugation step (17,600xg, 18 min, 4°C). The pellets were washed with the isotonic SEM buffer (250 mM sucrose, 1 mM EDTA, 10 mM MOPS/KOH, pH 7.2) containing 2 mM PMSF and centrifuged for 12 min as in the previous step. The pellet obtained after the high-speed spin is the mitochondrial fraction. The pellet was resuspended in SEM buffer, snap-frozen in liquid N_2_, and stored at –80°C until further use.

To obtain pure mitochondrial fractions, the mitochondrial pellet obtained after differential centrifugation was mixed with 2-3 mL of SEM buffer containing 1 mM PMSF and loaded on a sucrose step gradient (20, 30, 40, 50, and 60% (w/v) sucrose in 10 mM MOPS/KOH pH 7.4, 100 mM KCl, 1 mM EDTA, 1 mM PMSF). The gradients were centrifuged (210,000 x g, 16 h, 4°C) using a swing-out rotor (SW40Ti). The two fractions between the 40% and 60% phases, that correspond to the mitochondrial fractions, were collected and diluted with 35 mL SEM buffer containing 2 mM PMSF. A pure mitochondrial pellet was harvested by centrifugation of the suspension (17600 x g, 12 min, 4°C). The pellet was resuspended in SEM buffer, snap-frozen, and stored at –80°C until further use.

### Subcellular fractionation

Spheroplasts were prepared as for the isolation of mitochondria and next resuspended in SCF homogenization buffer (0.6 M sorbitol, 1 mM EDTA, 2 mM PMSF, 0.04 % (w/v) fatty-acid free BSA, 10 mM Tris, pH 7.4) and dounced 15 times using a homogenization Douncer. Samples corresponding to whole cell lysates (WCL) were taken and the lysate was subjected to two clarifying spins (2000xg, 5 min, 4°C) to remove nuclei and cell debris in the pellet. The mitochondrial fraction was pelleted by centrifugation (17600xg, 18 min, 4 °C). To isolate ER/microsomes from cytosolic fractions, 20 mL of the supernatant (post-mitochondrial fraction, PMF) were transferred to ultracentrifugation tubes and centrifuged (200000xg, 1 h, 4°C) using a Ti60 fixed angle rotor. The supernatant represents the cytosolic fraction. The pellet (consisting of ER/microsomes) was resuspended in 2 mL SEM buffer (with 2 mM PMSF), homogenized using a dounce homogenizer, and aliquoted. All samples (WCL, mitochondria, PMF, cytosolic fraction, and ER/microsomes) were snap-frozen and stored at –80 °C until further use.

### Isolation of lipid droplets

LDs were isolated according to a modified protocol (Zinser and Daum, 1995). Spheroplasts were resuspended in 25 mL LD buffer C (10 mM MES-Tris pH 6.9, 12 % (w/w) Ficoll 400, 0.2 mM EDTA, 2 mM PMSF) and homogenized 15x on ice using a Douncer with a loose-fitting pestle. Cell debris was removed by centrifugation (twice 2000xg, 5 min, 4°C) and the lysate was centrifuged (17600xg, 18 min, 4°C). The supernatant was transferred to ultracentrifuge tubes, mixed 1:1 with LD buffer C, and centrifuged (100000xg, 1 h, 4°C, SW40 rotor). The floating layer was then mixed with LD buffer C (1:1), treated with a homogenization Douncer (10 times), transferred to a fresh ultracentrifuge tube, overlayed with LD buffer D (10 mM MES-Tris pH 6.9, 8 % (w/w) Ficoll 400, 0.2 mM EDTA, 2 mM PMSF, 12 mL total volume) and centrifuged (100000xg, 30 min, 4°C, SW40 rotor). The floating layer was gently resuspended with a homogenization Douncer (10 times) in LD buffer E (10 mM MES-Tris pH 6.9, 8 % (w/w) Ficoll 400, 0.6 mM sorbitol, 0.2 mM EDTA, 2 mM PMSF), transferred to a fresh ultracentrifuge tube and overlayed with LD buffer F (10 mM MES-Tris pH 6.9, 0.25 mM sorbitol, 0.2 mM EDTA, 2 mM PMSF) to a final volume of 12 mL. After centrifugation (100000xg, 30 min, 4°C, SW40 rotor), the floating layer was collected, homogenized and aliquoted. Aliquots were snap-frozen and stored at –80 °C until further use.

### Isolation of vacuoles

Vacuoles were isolated according to a published protocol with some modifications (Kagohashi et al., 2023). Harvested spheroplasts were resuspended in 25 mL VC buffer A (10 mM MES-Tris, pH 6.9, 0.2 M sorbitol, 12% (w/v) Ficoll 400, 0.1 mM MgCl_2_) and homogenized using a loose-fitting pestle by douncing (15 times). The lysate was transferred to a fresh ultracentrifuge tube, layered with 10 mL VC buffer B (10 mM MES-Tris, pH 6.9, 0.2 M sorbitol, 8% (w/v) Ficoll 400, 0.1 mM MgCl_2_) and centrifuged (72000xg, 30 min, 4°C, SW28 rotor). The pellet was further processed to reisolate mitochondrial fractions as described below. The floating layer was mixed with 9 mL VC buffer B, treated with a homogenization Douncer (10 times), transferred to a fresh ultracentrifuge tube and overlayed with 10 mL VC buffer B’ (10 mM MES-Tris, pH 6.9, 0.2 M sorbitol, 4% (w/v) Ficoll 400, 0.1 mM MgCl_2_) and 10 mL VC buffer C (10 mM MES-Tris, pH 6.9, 0.2 M sorbitol, 0.1 mM MgCl_2_). After centrifugation (100000xg, 30 min, 4°C, SW28 rotor), the band at the VC buffer B’-VC buffer C interface (0-4% Ficoll) corresponding to the vacuolar fraction was collected with a Pasteur pipette, treated with a homogenization Douncer (10 times), aliquoted, snap-frozen and stored at –80 °C until further use.

To reisolate mitochondria after isolating vacuoles, the pellet resulting from the first centrifugation step was resuspended in 30 mL SEM buffer with 2 mM PMSF and the suspension was clarified by two centrifugation steps (2000xg, 5 min, 4°C). Next, the supernatant was centrifuged (17600xg, 18 min, 4°C). The resulting pellet containing the mitochondrial fraction was finally aliquoted, snap-frozen in liquid nitrogen and stored at –80 °C.

### Alkaline extraction

Crude mitochondrial fractions (100 µg) were transferred to an ultracentrifuge tube and centrifuged (17600xg, 12 min, 2°C). The pellet was resuspended in 100 µL 20 mM HEPES/KOH (pH 7.5) and 100 µL of freshly prepared 200 mM Na_2_CO_3_ solution was added. After incubation on ice for 30 min, the samples were centrifuged (100000xg, 30 min, 2°C). The pellet was resuspended in sample buffer. The supernatant was subjected to trichloroacetic acid (TCA) precipitation, and the resulting pellet was also resuspended in sample buffer. The samples were heated at 95°C for 10 min and analyzed by SDS-PAGE and immunodecoration. All antibodies used in this study are listed in Supplementary Table S7.

### Preparation of whole cell lysates by alkaline lysis

Pellets of cells corresponding to a total OD_600_ of 5 were harvested (3000xg, 5 min, RT), washed with water, and resuspended in 400 µL of 0.1 M NaOH. After incubation for 5 min at RT, cell pellets were harvested (3000xg, 5 min, RT), resuspended in 200 µL sample buffer, and heated for 10 min at 95 °C. Of each sample, 40 µL were analyzed by SDS-PAGE.

### Protease protection assay

Crude mitochondria (50 µg) were resuspended in 1 mL of either SEM buffer or 20 mM HEPES/KOH (pH 7.5) in the presence/absence of 1 % (v/v) Triton-X100. Proteinase K (PK) was added to a final concentration of 50 µg/mL and the samples were incubated for 30 min on ice. Organelles in SEM without PK served as an input control. The reactions were stopped by adding PMSF (2 mM final concentration) followed by incubation for 10 min on ice. All samples were subjected to TCA precipitation, resuspended in 40 µL sample buffer, heated at 95 °C for 10 min, and subjected to SDS-PAGE and immunodecoration.

### Co-Immunoprecipitation

Pellets of 50 mL mid-logarithmic yeast cells were prepared by centrifugation (3000xg, 5 min, 4°C), washed with ice-cold water, and resuspended in 1 mL IP buffer (10 mM Tris, 300 mM NaCl, 1 % (v/v) Tx-100, pH 7.5; 2 mM PMSF and 1 tablet of protease inhibitor cocktail (Merck) per 20 mL). Glass beads (Ø 0.5 mm, 600 mg) were added to the suspension, and the mixture was vortexed four times for 30 s with 30 s cooling breaks on ice in between. Next, the glass beads and cellular debris were removed (2000xg, 2 min, 4°C). The supernatant was transferred to the pre-washed anti-HA magnetic beads (washed thrice with 1 mL IP buffer) and incubated overnight on an overhead rotor (12 rpm, 4 °C). Next, the beads were collected with a magnetic rack and washed three times as described above with 1 mL IP buffer. Finally, collected magnetic beads were resuspended in 50 µL SDS sample buffer and heated for 10 min at 95 °C (IP sample). IP samples were further processed and subjected to mass spectrometric analysis (Proteome Center Tübingen) or analyzed by western blotting. To the latter end, 20 % of the input, 20 % of the unbound fraction and the complete elution fraction were mixed with sample buffer, heated for 10 min at 95 °C, and subjected to SDS-PAGE and Western blot analysis.

### BN-PAGE

Crude mitochondria (100 µg) were solubilized in 100 µL of BN solubilization buffer (20 mM Tris, 0.1 mM EDTA, 50 mM NaCl, 10 % (v/v) glycerol, pH 7.4, 2 mM PMSF) supplemented with Triton X-100 (0.5% (v/v). The solution was incubated for 30 min on ice and subjected to a clarifying spin (20000xg, 30 min, 4°C). The supernatant was mixed with the loading dye (5% (w/v) Coomassie blue G, 500 mM 6-amino-N-caproic acid, 100 mM Bis-Tris, pH 7.0) and analyzed on a 4-8% acrylamide blue native gel. The gels were blotted onto a PVDF membrane and further analyzed by immunodecoration.

### Fluorescence microscopy

For visualization of different organelles and subcellular structures *in vivo*, yeast cells were analyzed by confocal fluorescence microscopy. The Spinning disk microscope Zeiss Axio Examiner Z1 equipped with a CSU-X1 real-time confocal system and SPOT Flex charge-coupled device camera was used. Samples were observed using Objective Plan-Apochromat 63x / 1.4 Oil DIC M27 at room temperature. Images were acquired using the VisiView software and analyzed using ImageJ (Fiji). Z-stacks were taken in 0.2 µm step size.

### Flow cytometry analysis for quantification of lipid droplets

To quantify relative amounts of lipid droplets in cells, fluorescence intensity of endogenously labelled Pet10-mNG was analyzed by flow cytometry. The measurements were performed using a CytoFLEX S V0-B2-Y4-R0 flow cytometer with the CytExpert acquisition software. The cell suspension was analyzed with a flow rate of 60 µL/min and recorded until 1,000,000 cells (events) were counted. The mean FITC area (with standard deviation) was calculated for 1,000,000 cells of 3 independent experiments and plotted as mean fluorescence.

### Mass spectrometry

#### Proteomic sample preparation

Control (WT) cells, cells lacking peroxisomes (Δpex), and cells lacking the four enzymes required for triacylglycerol (TAG) and sterol ester (SE) synthesis (ΔLD cells) were grown in SILAC medium containing light, medium, or heavy labeled lysine, respectively. In all strains *LYS1* was deleted.

Crude mitochondrial fractions were isolated as described above from three independent biological replicates. Proteins were extracted and reduced with 1 mM dithiothreitol (DTT) for 1 h at room temperature while shaking. Alkylation was performed with 5.5 mM iodoacetamide (IAA) for 1 h at room temperature in the dark. Proteins were pre-digested with LysC (Wako Chemicals) for 3 h at room temperature. Subsequently, four volumes of water and additional LysC were added, and digestion continued overnight. Reactions were stopped by acidification to approximately 0.1% (v/v) trifluoroacetic acid (TFA).

Vlf1-3xHA and control samples were analyzed in three independent biological replicates. Following immunoprecipitation, proteins were separated by SDS-PAGE using 4–12% NuPAGE Bis-Tris gels (Invitrogen) for 7 min at 200 V and visualized by Coomassie staining. Gel regions containing the protein samples were excised and subjected to in-gel tryptic digestion as described previously (Shevchenko et al., 2006). Peptides were desalted and purified using C18 StageTips (Empore). C18 disks were activated with methanol and equilibrated with Solvent A* (2% (v/v) acetonitrile, 1% (v/v) formic acid). Samples were loaded onto the StageTips, washed with Solvent A (0.1% (v/v) formic acid), and eluted with 50 µL Solvent B (80% (v/v) acetonitrile, 0.1% (v/v) formic acid). Acetonitrile was removed by vacuum centrifugation, and samples were adjusted to the appropriate loading volume using Solvent A and 10% (v/v) Solvent A*.

#### Liquid chromatography-MS sample preparation and measurement

Peptides were analyzed on an Exploris 480 mass spectrometer (Thermo Fisher Scientific) coupled to a VanquishNeo UHPLC system (Thermo Fisher Scientific). Chromatographic separation was performed on a 20 cm × 75 µm analytical column packed in-house with ReproSil-Pur C18-AQ 1.9 µm resin (Dr. Maisch GmbH). Peptides were loaded at 40°C with a flow rate of 1 µL/min under a maximum backpressure of 1000 bar, and eluted using a 60-min gradient from 4% to 95% Solvent B in Solvent A at a constant flow rate of 200 nl/min.

Mass spectrometric analysis was performed in positive ion mode using data-dependent acquisition. Full MS scans were acquired over an m/z range of 300–1750 at a resolution of 60,000. The 20 most abundant multiply charged precursor ions were selected for HCD fragmentation with a dynamic exclusion time of 30 s. MS/MS spectra were acquired at a resolution of 15,000. The automatic gain control (AGC) was set to custom, with a normalized AGC target of 50% and an absolute value of 5.000e-4, and the maximum injection time was set to auto.

#### Mass spectrometry data processing

Raw acquired files were processed using MaxQuant software (version 2.2.0.0.) and searched against the Uniprot *Saccharomyces cerevisiae* database (6,091 entries, downloaded 2024/01/30) and commonly observed protein contaminants. For MS and MS/MS, the peptide mass tolerance was set at 4.5 ppm and 20 ppm, respectively.

For the SILAC-based mitochondrial proteomics dataset, LysC was specified as the proteolytic enzyme, whereas trypsin was used for the Vlf1-3xHA immunoprecipitation samples. Up to two missed cleavages were allowed. Carbamidomethylation of cysteine residues was set as a fixed modification, while methionine oxidation and protein N-terminal acetylation were included as variable modifications.

The false discovery rate was set to 1% at both the peptide and protein levels. For label-free quantification, a minimum ratio count of two was requested. Intensity-based absolute quantification (iBAQ) was enabled. All other parameters remained at their default settings.

#### Mass spectrometry data analysis

Downstream analysis of MaxQuant output tables was performed using Perseus (version 2.0.10.0). Potential contaminants, reverse database hits, and proteins identified only by site were filtered out prior to analysis. Proteins were functionally annotated using Gene Ontology (GO) Biological Process, Cellular Component, and Molecular Function terms, Kyoto Encyclopedia of Genes and Genomes (KEGG) annotations, and the Saccharomyces Genome Database (SGD) mitochondrial protein database.

For the SILAC-based mitochondrial proteomics dataset, the summed intensities of the light/medium, light/heavy, and heavy/medium channels were log10-transformed. Normalized SILAC ratios were log2-transformed. Rows were filtered for at least two valid values per protein. Median protein ratios and summed intensity were calculated. Volcano plots displaying log10 summed intensity versus log2 SILAC ratio were generated. Proteins exhibiting changes greater than ±0.5 on the log2 scale were considered differentially abundant. Mitochondrial annotated proteins of unknown function were selected for highlighted.

For the Vlf1-3xHA immunoprecipitation dataset, protein intensities were log10-transformed and filtered for at least two valid values across all samples. Missing values were imputed in Perseus using a normal distribution (width = 0.3, downshift = 1.8) separately for each sample column. A Student’s t-test was performed to identify significantly enriched proteins. Volcano plots were generated using the t-test difference on the x-axis and the corresponding –log10 p-value on the y-axis.

Additional graphical visualization was performed in the R environment (version 4.1.1) and GraphPad (version 8.0.1), while figures were edited using Adobe Illustrator.

### High content visual screen

Yeast cells were visualized in synthetic media used containing 6.7 g/L yeast nitrogen base with ammonium sulfate and 2% glucose (SD) supplemented with a complete amino acid mix (oMM composition). The glucose and nitrogen starvation medium (S-N) used in this study consisted of 1.7 g/L yeast nitrogen base without ammonium sulfate. Diploid cells were selected using antibiotics at the following concentrations: 500 mg/L Geneticin and 200 mg/L Nourseothricin. To generate the diploid library used in this study, we utilized an automated mating method (described in (Cohen and Schuldiner, 2011)) using a RoToR bench-top colony arraying instrument (Singer Instruments). The mated strain was yMS9201 (Vlf1 tagged with mNG at the C-terminus), and the library of strains was taken from the Tef2-mCherry collection (Weill et al., 2018). The library and the query strain were mated on YPD agar plates, then transferred onto YPD + hygromycin + clonNAT plates for diploid selection.

Automated microscopy was used to screen cells. The library was transferred from 384-well agar plates to 384-well polystyrene plates containing 100 μL of selection medium (SD complete (oMM) + hygromycin + clonNAT) and incubated overnight in a Liconic incubator at 30°C. Cells were then back-diluted into 384-well plates containing 95 μL of S-N (glucose and nitrogen starvation medium). After 6 hours, 50 μL from each well were transferred to a glass-bottomed 384-well microscopy plate coated with Concanavalin A. Following a 20-minute incubation at 25°C, the plates were washed once with S-N medium and imaged. The screen was imaged using a 60× air objective (NA 0.9) and an ORCA-Flash4.0 digital camera (Hamamatsu). Images were recorded using 488 nm and 561 nm laser illumination for the green and red channels, respectively. All steps in the automated imaging process were performed using a Freedom EVO liquid handler (Tecan). Microscopy images were cropped, colored, slightly adjusted for brightness/contrast, and analyzed using ImageJ (Schneider et al., 2012).

## Acknowledgements

We thank E. Kracker for technical assistance. We are grateful to C. Ungermann, T. Becker and R. Lill for the kind gift of antibodies. We also would like to thank J. Ewald for providing several plasmids. This work was supported by the Deutsche Forschungsgemeinschaft through Research Training Group 2364/2 (Z.F., C.C.R., K.S.D., B.M., and D.R.). Work in the group of K.S. Dimmer was supported by the Wilhelm Schuler Stiftung. We are grateful to Y. Asraf for her help with the high content screens. The robotic system of the Schuldiner lab was purchased through the kind support of the Blythe Brenden-Mann Foundation. MS is an Incumbent of the Dr. Gilbert Omenn and Martha Darling Professorial Chair in Molecular Genetics.

## Author Contributions

ZF, CCR, BRG, SR, PD, OB and KSD performed experiments and analyzed data together with MS, BM and DR. KSD designed the study and wrote the manuscript with major input from ZF and contributions from MS, BM and DR.

## Figure Legends

**Supplementary Figure S1:**
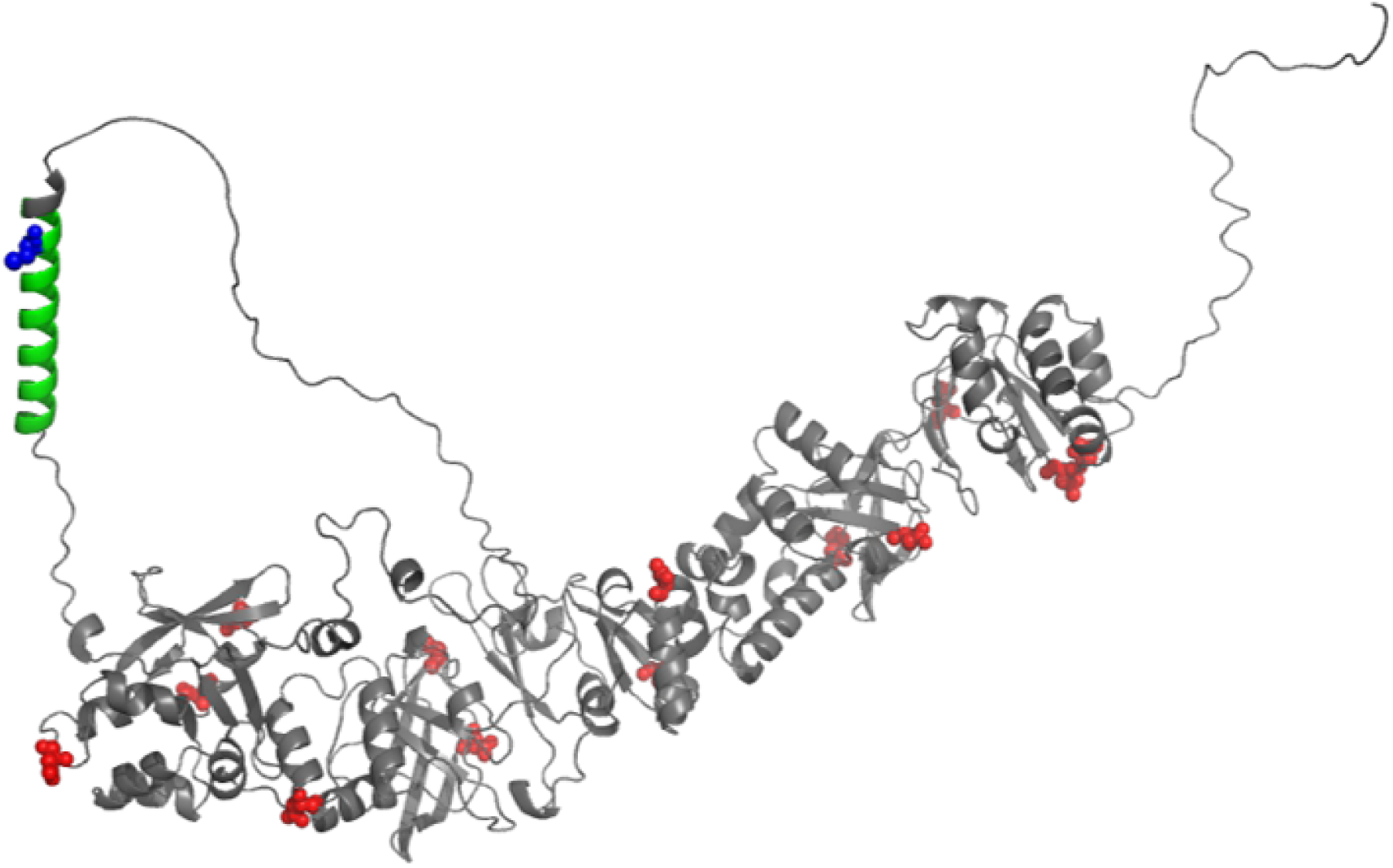
Predicted structure of Vlf1. The structure of Vlf1 was predicted with AlphaFold 2. The protein contains a putative transmembrane domain (highlighted in green) with a predicted palmytoylation site (C780, highlighted in blue). The predicted structure shows also potential glycosylation sites (highlighted in red).

**Supplementary Figure S2:**
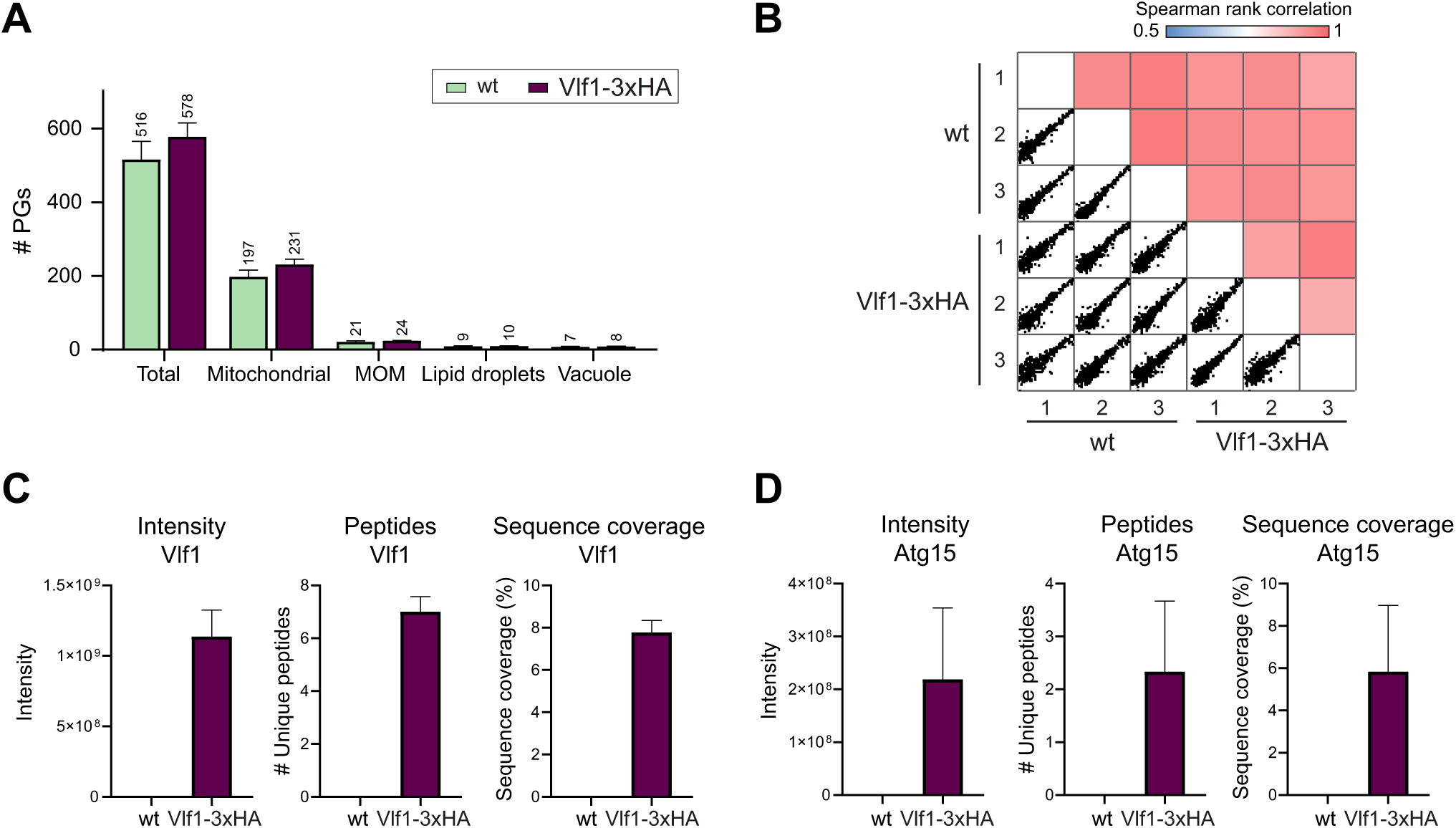
Quality evaluation of the coimmunprecipitation of Vlf1-3xHA. **(A)** Number and subcellular annotation of proteins identified in the wt control and Vlf1-3xHA immunoprecipitation samples. A total of 516 proteins were identified in WT samples and 579 proteins in Vlf1-3xHA immunoprecipitates. **(B)** Pearson correlation analysis of three independent biological replicates demonstrating high reproducibility of the proteomic dataset (r > 0.88). **(C)** Detection of Vlf1 in the proteomics dataset. Shown are protein intensity, number of identified peptides, and sequence coverage for Vlf1 in WT control and Vlf1-3xHA immunoprecipitation samples. **(D)** Detection of Atg15, the main interacting protein, in the proteomics dataset. Shown are protein intensity, number of identified peptides, and sequence coverage for Atg15 in WT control and Vlf1-3xHA immunoprecipitation samples.

**Supplementary Figure S3:**
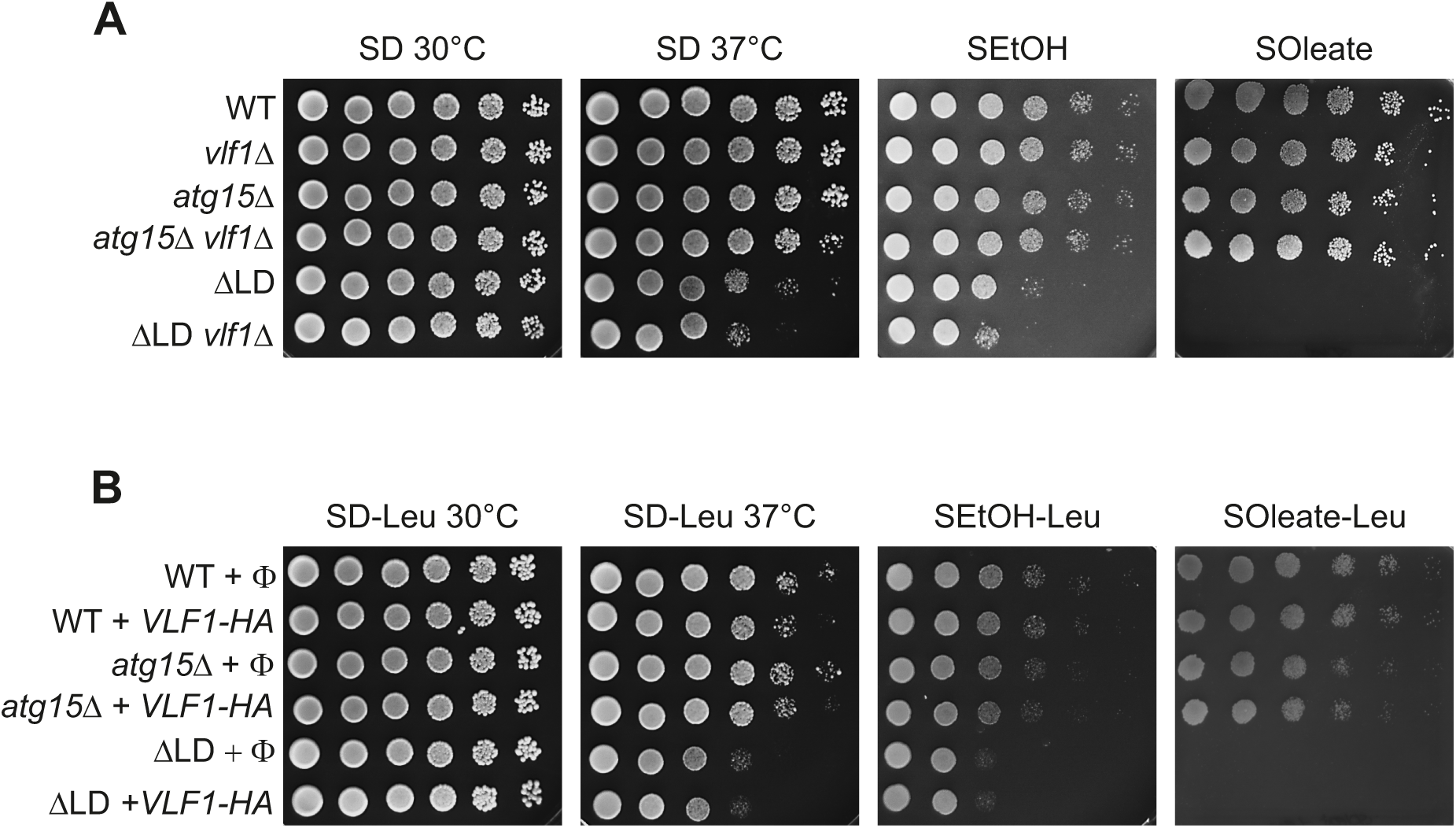
Deletion or overexpression of *VLF1* has no effect on the growth of yeast cells. The growth of cells lacking *VLF1* (A) or overexpressing *VLF1* from a plasmid (B) was analyzed by drop dilution assay of 1:5 serial dilutions on synthetic medium containing glucose (SD) at either 30 °C or 37 °C, or containing ethanol (SEtOH) or oleate (SOleate) at 30 °C. The growth of cells lacking or overexpressing Vlf1 was also analyzed for cells that harbored also the deletion of *ATG15* (*atg15*D) or completely lacked lipid droplets (DLD).

**Supplementary Table S2:**
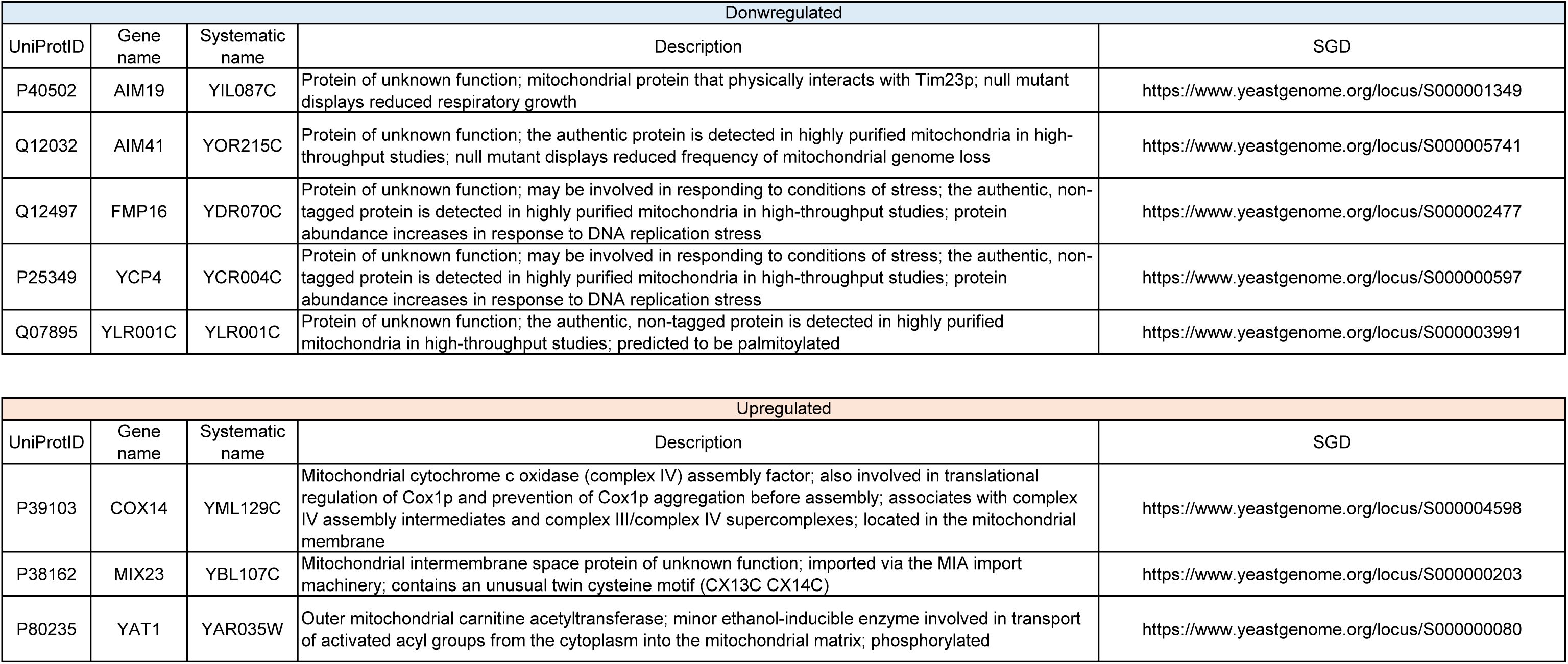
Mitochondrial proteins with altered expression in ΔLD yeast cells.

| Downregulated |  |  |  |  |
| --- | --- | --- | --- | --- |
| UniProtID | Gene name | Systematic name | Description | SGD |
| P40502 | AIM19 | YIL087C | Protein of unknown function; mitochondrial protein that physically interacts with Tim23p; null mutant displays reduced respiratory growth | https://www.yeastgenome.org/locus/S000001349 |
| Q12032 | AIM41 | YOR215C | Protein of unknown function; the authentic protein is detected in highly purified mitochondria in high-throughput studies; null mutant displays reduced frequency of mitochondrial genome loss | https://www.yeastgenome.org/locus/S000005741 |
| Q12497 | FMP16 | YDR070C | Protein of unknown function; may be involved in responding to conditions of stress; the authentic, non-tagged protein is detected in highly purified mitochondria in high-throughput studies; protein abundance increases in response to DNA replication stress | https://www.yeastgenome.org/locus/S000002477 |
| P25349 | YCP4 | YCR004C | Protein of unknown function; may be involved in responding to conditions of stress; the authentic, non-tagged protein is detected in highly purified mitochondria in high-throughput studies; protein abundance increases in response to DNA replication stress | https://www.yeastgenome.org/locus/S000000597 |
| Q07895 | YLR001C | YLR001C | Protein of unknown function; the authentic, non-tagged protein is detected in highly purified mitochondria in high-throughput studies; predicted to be palmitoylated | https://www.yeastgenome.org/locus/S000003991 |

| Upregulated |  |  |  |  |
| --- | --- | --- | --- | --- |
| UniProtID | Gene name | Systematic name | Description | SGD |
| P39103 | COX14 | YML129C | Mitochondrial cytochrome c oxidase (complex IV) assembly factor; also involved in translational regulation of Cox1p and prevention of Cox1p aggregation before assembly; associates with complex IV assembly intermediates and complex III/complex IV supercomplexes; located in the mitochondrial membrane | https://www.yeastgenome.org/locus/S000004598 |
| P38162 | MIX23 | YBL107C | Mitochondrial intermembrane space protein of unknown function; imported via the MIA import machinery; contains an unusual twin cysteine motif (CX13C CX14C) | https://www.yeastgenome.org/locus/S000000203 |
| P80235 | YAT1 | YAR035W | Outer mitochondrial carnitine acetyltransferase; minor ethanol-inducible enzyme involved in transport of activated acyl groups from the cytoplasm into the mitochondrial matrix; phosphorylated | https://www.yeastgenome.org/locus/S000000080 |

**Supplementary Table S3:** Proteins tagged with mCherry forming punctate structures colocalizing with Vlf1-mNG.

| Supplementary Table S3: Proteins tagged with mCherry forming punctate structures colocalizing with Vif1-mNG |  |  |  |
| --- | --- | --- | --- |
| Systematic Name | Gene Name | Function according to SGD | Organelle |
| YIR034C | LYS1 | Saccharopine dehydrogenase (NAD<sup>+</sup>, L-lysine-forming); catalyzes the conversion of saccharopine to L-lysine, which is the final step in the lysine biosynthesis pathway; also has mRNA binding activity | cytosol |
| YNL048W | ALG11 | Alpha-1,2-mannosyltransferase; catalyzes sequential addition of the two terminal alpha 1,2-mannose residues to the Man5GlcNAc2-PP-dolichol intermediate during asparagine-linked glycosylation in the ER | ER |
| YKL179C | COY1 | Golgi membrane protein with similarity to mammalian CASP; genetic interactions with GOS1 (encoding a Golgi snare protein) suggest a role in Golgi function | golgi |
| YMR313C | TGL3 | Bifunctional triacylglycerol lipase and LPE acyltransferase; major lipid particle-localized triacylglycerol (TAG) lipase; catalyzes acylation of lysophosphatidylethanolamine (LPE), a function which is essential for sporulation; protein level and stability of Tgl3p are markedly reduced in the absence of lipid droplets; required with Tgl4p for timely bud formation | LD |
| YMR148W | OSW5 | Protein of unknown function with possible role in spore wall assembly; predicted to contain an N-terminal transmembrane domain; osw5 null mutant spores exhibit increased spore wall permeability and sensitivity to beta-glucanase digestion | LD |
| YOL047C | LDS2 | Protein Involved in spore wall assembly; localizes to lipid droplets found on or outside of the prospore membrane shares similarity with Lds1p and Rrt8p, and a strain mutant for all 3 genes exhibits reduced dityrosine fluorescence relative to the single mutants; green fluorescent protein (GFP)-fusion protein localizes to the cytoplasm in a punctate pattern | LD |
| YOR081C | TGL5 | Bifunctional triacylglycerol lipase and LPA acyltransferase; lipid particle-localized triacylglycerol (TAG) lipase involved in triacylglycerol mobilization; catalyzes acylation of lysophosphatidic acid (LPA); potential Cdc28p substrate; TGL5 has a paralog, TGL4, that arose from the whole genome duplication | LD |
| YOL048C | RRT8 | Protein involved in spore wall assembly; shares similarity with Lds1p and Lds2p and a strain mutant for all 3 genes exhibits reduced dityrosine fluorescence relative to the single mutants; identified in a screen for mutants with increased levels of rDNA transcription; green fluorescent protein (GFP)-fusion protein localizes to lipid particles; protein abundance increases in response to DNA replication stress | LD / sporewall |
| YLR389C | STE23 | Metalloprotease; involved in N-terminal processing of pro-a-factor to mature form; expressed in both haploids and diploids; one of two yeast homologs of human insulin-degrading enzyme (hIDE); homolog Axl1p is also involved in processing of pro-a-factor | mitochondrion |
| YNL115C | YNL115C | Putative protein of unknown function; green fluorescent protein (GFP)-fusion protein localizes to mitochondria YNL115C is not an essential gene | mitochondrion |
| YER015W | FAA2 | Medium chain fatty acyl-CoA synthetase; activates imported fatty acids; accepts a wide range of fatty acid chain lengths with a preference for medium chains, C9:0-C13:0; localized to the peroxisome; comparative analysis suggests that a mitochondrially targeted form may result from translation starting at a non-canonical codon upstream of the annotated start codon | mitochondrion / peroxisome |
| YKL057C | NUP120 | Subunit of the Nup84p subcomplex of the nuclear pore complex (NPC); contributes to nucleocytoplasmic transport and NPC biogenesis and is involved in establishment of a normal nucleocytoplasmic concentration gradient of the GTPase Gsp1p; also plays roles in several processes that may require localization of genes or chromosomes at the nuclear periphery, including double-strand break repair, transcription and chromatin silencing; homologous to human NUP160 | nucleus |
| YDR096W | GIS1 | Histone demethylase and transcription factor; regulates genes during nutrient limitation; activity modulated by proteasome-mediated proteolysis; has JmjC and JmjN domain in N-terminus that interact, promoting stability and proper transcriptional activity; contains two transactivating domains downstream of Jmj domains and a C-terminal DNA binding domain; relocates to the cytosol in response to hypoxia; GIS1 has a paralog, RPH1, that arose from the whole genome duplication | nucleus / cytosol |
| YKR009C | FOX2 | 3-hydroxyacyl-CoA dehydrogenase and enoyl-CoA hydratase; multifunctional enzyme of the peroxisomal fatty acid beta-oxidation pathway; mutation is functionally complemented by human HSD17B4 | peroxisome |
| YLR027C | AAT2 | Cytosolic aspartate aminotransferase involved in nitrogen metabolism; localizes to peroxisomes in oleate-grown cells | peroxisome |
| YOL044W | PEX15 | Tail-anchored type II integral peroxisomal membrane protein; required for peroxisome biogenesis; cells lacking Pex15p mislocalize peroxisomal matrix proteins to cytosol; overexpression results in impaired peroxisome assembly | peroxisome |
| YPR128C | ANT1 | Peroxisomal adenine nucleotide transporter; involved in beta-oxidation of medium-chain fatty acid; required for peroxisome proliferation | peroxisome |
| YKR001C | VPS1 | Dynamin-like GTPase required for vacuolar sorting; also involved in actin cytoskeleton organization, endocytosis, late Golgi-retention of some proteins, regulation of peroxisome biogenesis | vacuole |
| YKL103C | APE1 | Vacuolar aminopeptidase ysc1; zinc metalloproteinase that belongs to the peptidase family M18; often used as a marker protein in studies of autophagy and cytosol to vacuole targeting (CVT) pathway; protein increases in abundance and relative distribution to cytoplasmic foci increases upon DNA replication stress | vacuole |
| YHR113W | APE4 | Cytoplasmic aspartyl aminopeptidase with possible vacuole function; Cvt pathway cargo protein; cleaves unblocked N-terminal acidic amino acids from peptide substrates; forms a 12-subunit homo-oligomer; M18 metalloprotease family | vacuole |
| YAL002W | VPS8 | Membrane-binding component of the CORVET complex; involved in endosomal vesicle tethering and fusion in the endosome to vacuole protein targeting pathway; interacts with Vps21p; contains RING finger motif | vacuole |
| YDR229W | IVY1 | Phospholipid-binding protein that interacts with both Ypt7p and Vps33p; may partially counteract the action of Vps33p and vice versa, localizes to the rim of the vacuole as cells approach stationary phase | vacuole |

## Supplementary Tables S4-S7

**Supplementary Table S4:**
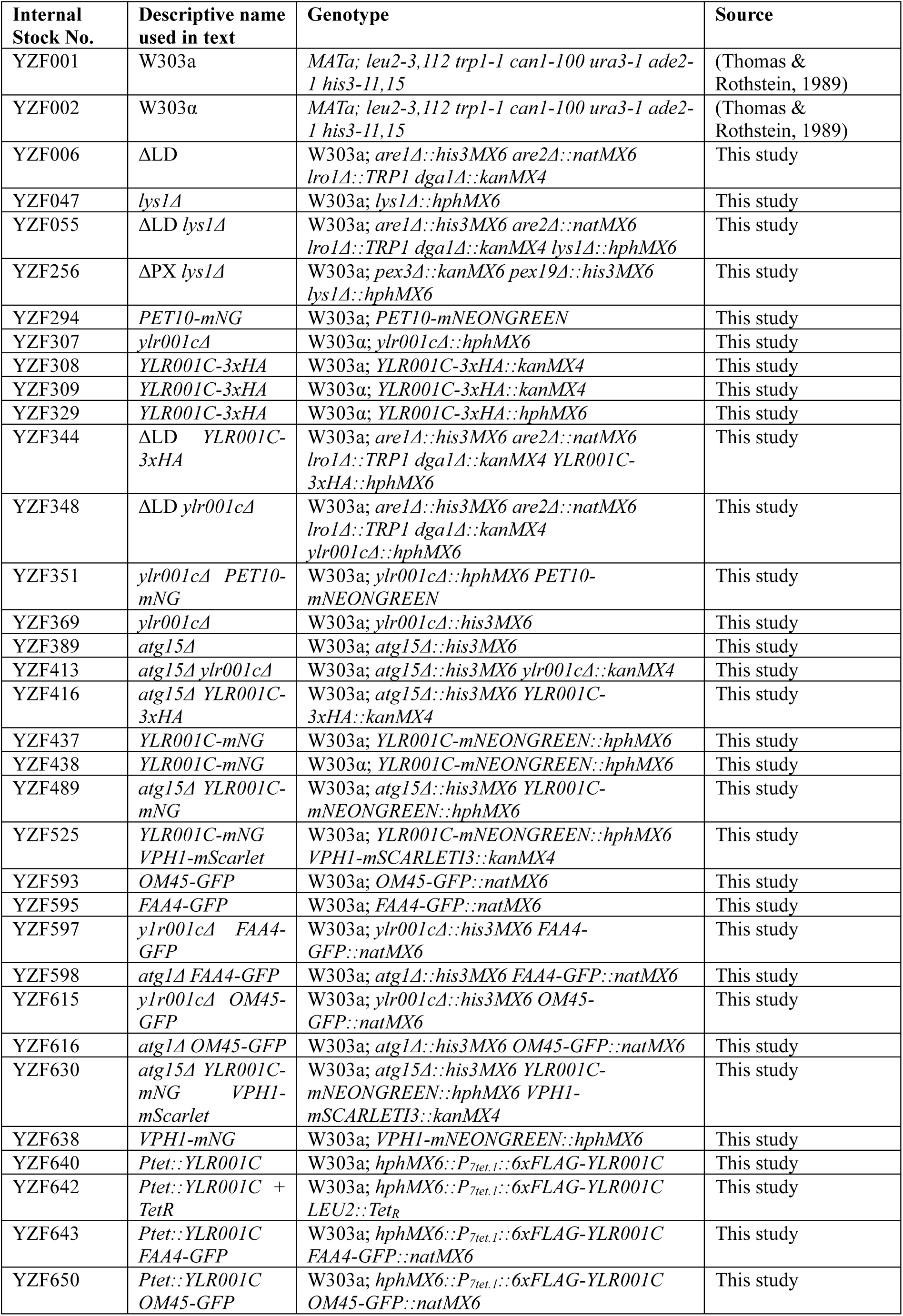
*S. cerevisiae* strains used in this study.

**Supplementary Table S5:**
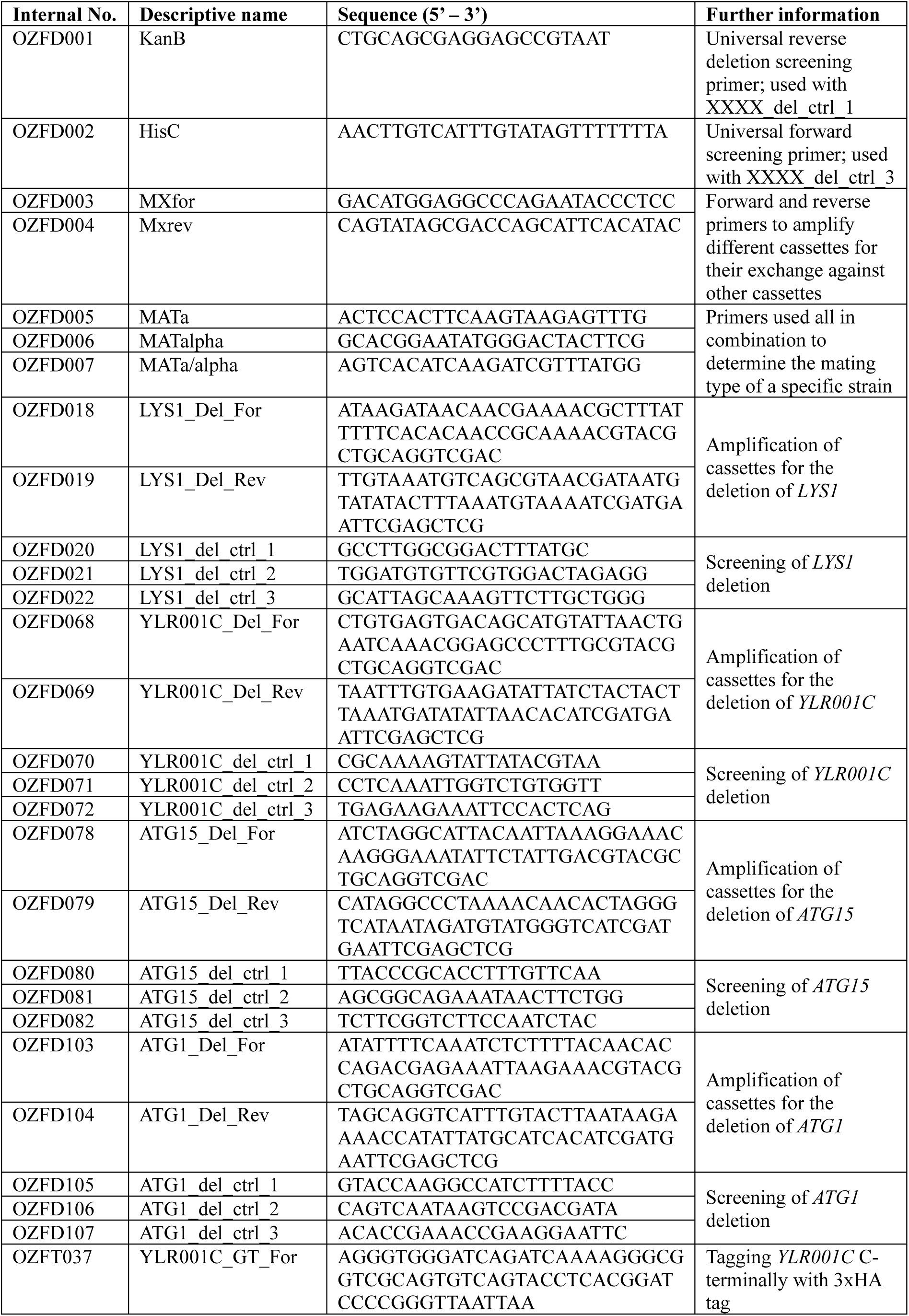
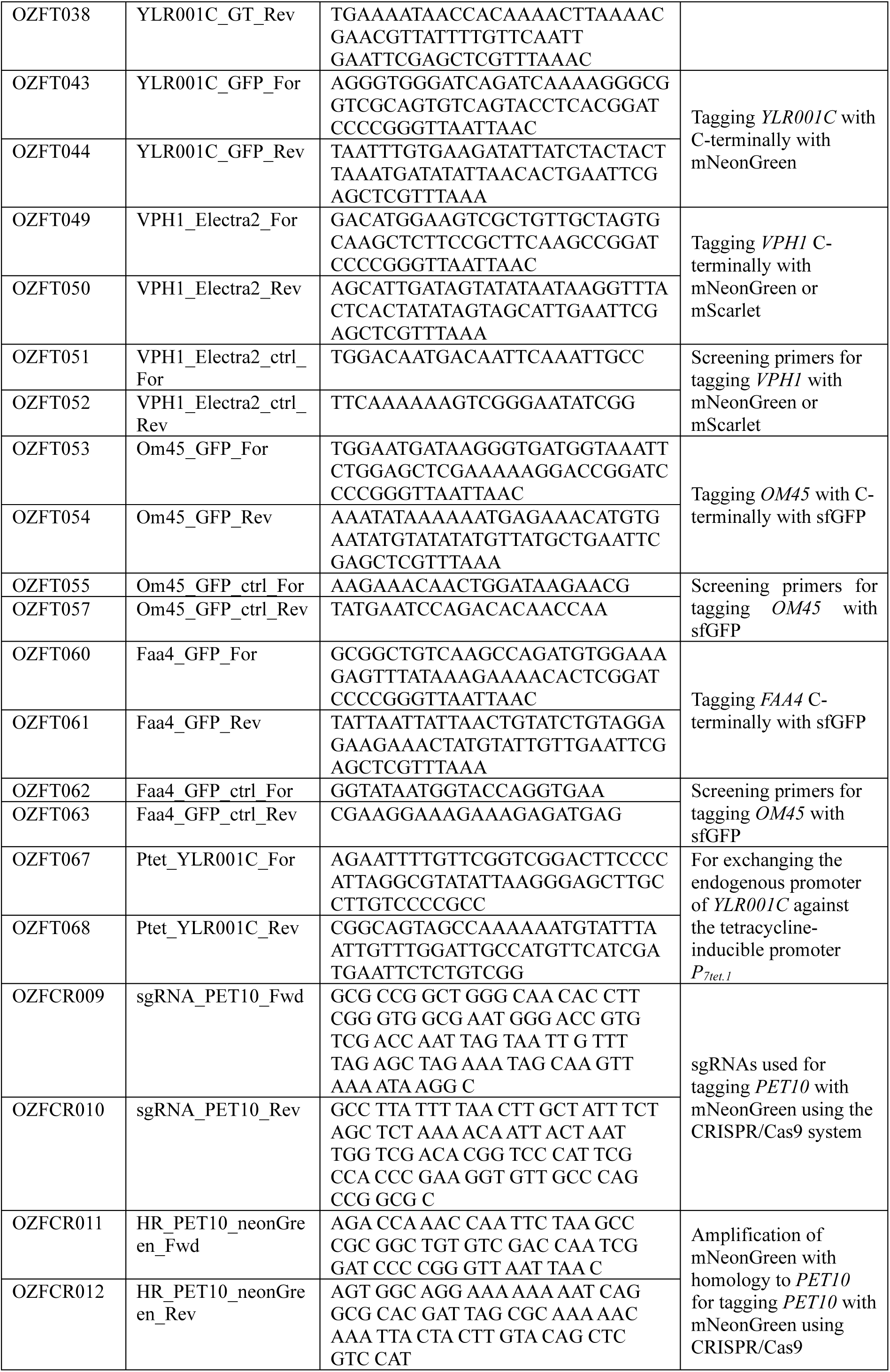
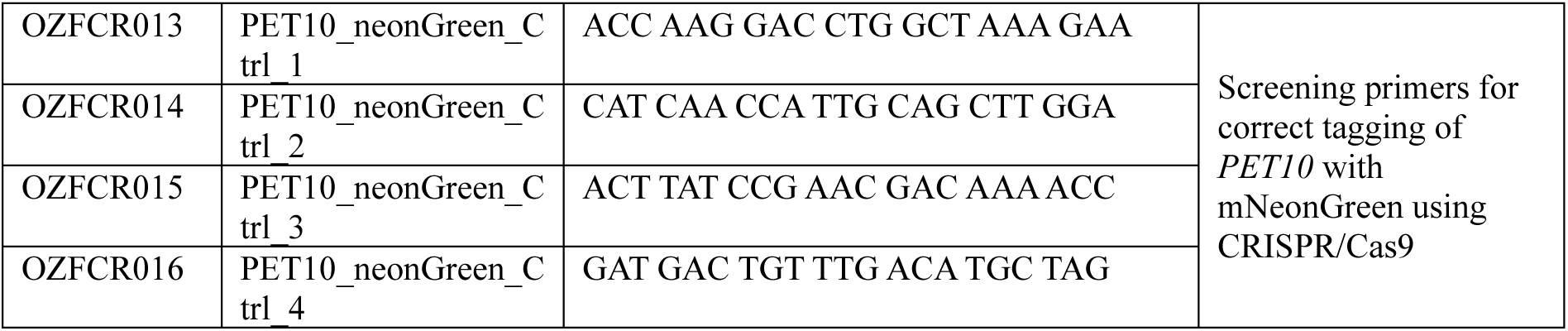
Oligonucleotides used in this study.

**Supplementary Table S6:** List of plasmids used in this study.

| Plasmid Number | Name | marker | Further information | Source |
| --- | --- | --- | --- | --- |
| pZF001 | pFA6a His3MX6 | Amp<sup>R</sup> | HIS3-deletion cassette | Lab Stock |
| pZF002 | pFA6a KanMX4 | Amp<sup>R</sup> | Kanamycin-deletion cassette | Lab Stock |
| pZF003 | pFA6a HphMX6 | Amp<sup>R</sup> | Hygromycin-deletion cassette | Lab Stock |
| pZF004 | pFA6a 3xHA-KanMX6 | Amp<sup>R</sup> | For C-terminal tagging with 3xHA | Lab Stock |
| pZF007 | pYX142 | Amp<sup>R</sup>,<br>*LEU2* | Yeast expression vector; empty plasmid | Lab Stock |
| pZF029 | Tet Repressor (Leu) | Amp<sup>R</sup>,<br>*LEU2* | Integrative tetracycline repressor; *AscI* digest for integration into the *LEU2* locus | Ewald Lab (pFS038) |
| pZF032 | Tet Promoter | Amp<sup>R</sup> | Tetracycline-inducible promoter | Ewald Lab (pFS042) |
| pZF041 | Cas9 | Amp<sup>R</sup> | Expresses Cas9 under the estradiol-inducible promoter | Ewald Lab (pFS046) |
| pZF045 | pFA6a_mNeonGreen-HphMX6 | Amp<sup>R</sup> | For C-terminal tagging with mNeonGreen | Ewald Lab (pYLB10) |
| pZF058 | pFA6a-sfGFP-NatMX6 | Amp<sup>R</sup> | For C-terminal tagging with sfGFP | Ewald Lab (pFS049) |
| pZF065 | pFA6a-mScarlet-I3-KanMX4 | Amp<sup>R</sup> | For C-terminal tagging with mScarlet | Schuldiner Lab (pMS1555) |
| pZF069 | pEX-A258-YLR001C | Amp<sup>R</sup> | Synthetic YLR001C; *EcoRI* and *XmaI* restriction sites for subcloning into yeast expression vector | Eurofins Genomics |
| pZF070 | pYX142_YLR001C-HA | Amp<sup>R</sup>,<br>*LEU2* | Expresses YLR001C-HA under the TPI promoter | This study |
| pZF074 | GFP-ATG8 | Amp<sup>R</sup>,<br>*URA3* | Expresses GFP-ATG8 under its endogenous promoter | Addgene (49425) |

**Supplementary Table S7:**
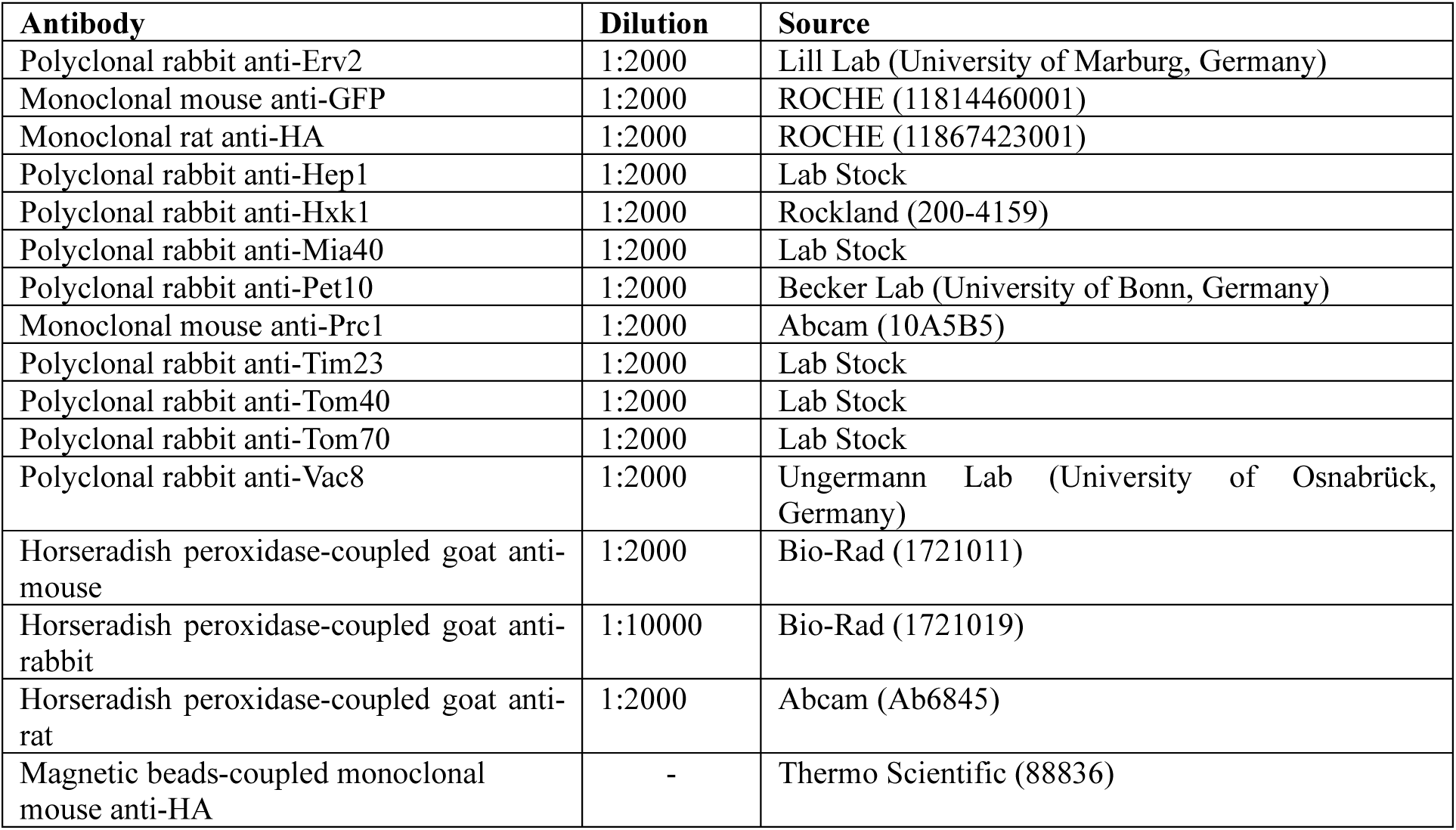
List of antibodies used in this study.

## References

1. Azizoglu, A., R. Brent, and F. Rudolf. 2021. A precisely adjustable, variation-suppressed eukaryotic transcriptional controller to enable genetic discovery. Elife. 10.

2. Baruch, D., I. Tsirkas, E. Sass, B. Dubreuil, Y. Asraf, A. Aharoni, M. Schuldiner, and O. Klein. 2025. Creation and validation of a proteome-wide yeast library for protein detection and analysis. J Cell Sci. 138.

3. Bischof, J., M. Salzmann, M.K. Streubel, J. Hasek, F. Geltinger, J. Duschl, N. Bresgen, P. Briza, D. Haskova, R. Lejskova, M. Sopjani, K. Richter, and M. Rinnerthaler. 2017. Clearing the outer mitochondrial membrane from harmful proteins via lipid droplets. Cell Death Discov. 3:17016.

4. Bohnert, M. 2020. Tether Me, Tether Me Not-Dynamic Organelle Contact Sites in Metabolic Rewiring. Dev Cell. 54:212–225.

5. Bohnert, M. 2026. I did it my way: the origins of lipid droplet heterogeneity. Curr Opin Cell Biol. 101:102665.

6. Cohen, Y., and M. Schuldiner. 2011. Advanced methods for high-throughput microscopy screening of genetically modified yeast libraries. Methods Mol Biol. 781:127–159.

7. Czabany, T., K. Athenstaedt, and G. Daum. 2007. Synthesis, storage and degradation of neutral lipids in yeast. Biochim Biophys Acta. 1771:299–309.

8. Daum, G., P.C. Bohni, and G. Schatz. 1982. Import of proteins into mitochondria. Cytochrome b2 and cytochrome c peroxidase are located in the intermembrane space of yeast mitochondria. J Biol Chem. 257:13028–13033.

9. Dimmer, K.S., and D. Rapaport. 2017. Mitochondrial contact sites as platforms for phospholipid exchange. Biochim Biophys Acta Mol Cell Biol Lipids. 1862:69–80.

10. Enkler, L., and A. Spang. 2024. Functional interplay of lipid droplets and mitochondria. FEBS Lett. 598:1235–1251.

11. Fairman, G., and M. Ouimet. 2022. Lipophagy pathways in yeast are controlled by their distinct modes of induction. Yeast. 39:429–439.

12. Garbarino, J., M. Padamsee, L. Wilcox, P.M. Oelkers, D. D’Ambrosio, K.V. Ruggles, N. Ramsey, O. Jabado, A. Turkish, and S.L. Sturley. 2009. Sterol and diacylglycerol acyltransferase deficiency triggers fatty acid-mediated cell death. J Biol Chem. 284:30994–31005.

13. Graef, M. 2018. Lipid droplet-mediated lipid and protein homeostasis in budding yeast. FEBS Lett. 592:1291–1303.

14. Henne, W.M. 2023. The (social) lives, deaths, and biophysical phases of lipid droplets. Curr Opin Cell Biol. 82:102178.

15. Henne, W.M., and S. Cohen. 2026. Heterogeneity, dynamics and organelle interactions of lipid droplets. Nat Rev Mol Cell Biol.

16. Hentges, P., B. Van Driessche, L. Tafforeau, J. Vandenhaute, and A.M. Carr. 2005. Three novel antibiotic marker cassettes for gene disruption and marker switching in Schizosaccharomyces pombe. Yeast. 22:1013–1019.

17. Herker, E., G. Vieyres, M. Beller, N. Krahmer, and M. Bohnert. 2021. Lipid Droplet Contact Sites in Health and Disease. Trends Cell Biol. 31:345–358.

18. Hiltunen, J.K., A.M. Mursula, H. Rottensteiner, R.K. Wierenga, A.J. Kastaniotis, and A. Gurvitz. 2003. The biochemistry of peroxisomal beta-oxidation in the yeast Saccharomyces cerevisiae. FEMS Microbiol Rev. 27:35–64.

19. Huh, W.K., J.V. Falvo, L.C. Gerke, A.S. Carroll, R.W. Howson, J.S. Weissman, and E.K. O’Shea. 2003. Global analysis of protein localization in budding yeast. Nature. 425:686–691.

20. Inokoshi, J., H. Tomoda, H. Hashimoto, A. Watanabe, H. Takeshima, and S. Omura. 1994. Cerulenin-resistant mutants of Saccharomyces cerevisiae with an altered fatty acid synthase gene. Mol Gen Genet. 244:90–96.

21. Janke, C., M.M. Magiera, N. Rathfelder, C. Taxis, S. Reber, H. Maekawa, A. Moreno-Borchart, G. Doenges, E. Schwob, E. Schiebel, and M. Knop. 2004. A versatile toolbox for PCR-based tagging of yeast genes: new fluorescent proteins, more markers and promoter substitution cassettes. Yeast. 21:947–962.

22. Jumper, J., R. Evans, A. Pritzel, T. Green, M. Figurnov, O. Ronneberger, K. Tunyasuvunakool, R. Bates, A. Zidek, A. Potapenko, A. Bridgland, C. Meyer, S.A.A. Kohl, A.J. Ballard, A. Cowie, B. Romera-Paredes, S. Nikolov, R. Jain, J. Adler, T. Back, S. Petersen, D. Reiman, E. Clancy, M. Zielinski, M. Steinegger, M. Pacholska, T. Berghammer, S. Bodenstein, D. Silver, O. Vinyals, A.W. Senior, K. Kavukcuoglu, P. Kohli, and D. Hassabis. 2021. Highly accurate protein structure prediction with AlphaFold. Nature. 596:583–589.

23. Kagohashi, Y., M. Sasaki, A.I. May, T. Kawamata, and Y. Ohsumi. 2023. The mechanism of Atg15-mediated membrane disruption in autophagy. J Cell Biol. 222.

24. Kotani, T., and H. Nakatogawa. 2026. Core principles of autophagy initiation mechanisms. Nat Struct Mol Biol. 33:408–419.

25. Kounakis, K., M. Chaniotakis, M. Markaki, and N. Tavernarakis. 2019. Emerging Roles of Lipophagy in Health and Disease. Front Cell Dev Biol. 7:185.

26. Marquardt, L., M. Montino, Y. Muhe, P. Schlotterhose, and M. Thumm. 2023. Topology and Function of the S. cerevisiae Autophagy Protein Atg15. Cells. 12.

27. Mathiowetz, A.J., and J.A. Olzmann. 2024. Lipid droplets and cellular lipid flux. Nat Cell Biol. 26:331–345.

28. Michaelis, A.C., A.D. Brunner, M. Zwiebel, F. Meier, M.T. Strauss, I. Bludau, and M. Mann. 2023. The social and structural architecture of the yeast protein interactome. Nature. 624:192–200.

29. Onishi, M., K. Yamano, M. Sato, N. Matsuda, and K. Okamoto. 2021. Molecular mechanisms and physiological functions of mitophagy. EMBO J. 40:e104705.

30. Pascual-Ahuir, A., S. Manzanares-Estreder, and M. Proft. 2017. Pro – and Antioxidant Functions of the Peroxisome-Mitochondria Connection and Its Impact on Aging and Disease. Oxid Med Cell Longev. 2017:9860841.

31. Petschnigg, J., H. Wolinski, D. Kolb, G. Zellnig, C.F. Kurat, K. Natter, and S.D. Kohlwein. 2009. Good fat, essential cellular requirements for triacylglycerol synthesis to maintain membrane homeostasis in yeast. J Biol Chem. 284:30981–30993.

32. Picca, A., J. Faitg, J. Auwerx, L. Ferrucci, and D. D’Amico. 2023. Mitophagy in human health, ageing and disease. Nat Metab. 5:2047–2061.

33. Reinders, J., R.P. Zahedi, N. Pfanner, C. Meisinger, and A. Sickmann. 2006. Toward the complete yeast mitochondrial proteome: multidimensional separation techniques for mitochondrial proteomics. J Proteome Res. 5:1543–1554.

34. Schneider, C.A., W.S. Rasband, and K.W. Eliceiri. 2012. NIH Image to ImageJ: 25 years of image analysis. Nat Methods. 9:671–675.

35. Schott, M.B., C.N. Rozeveld, S.G. Weller, and M.A. McNiven. 2022. Lipophagy at a glance. J Cell Sci. 135.

36. Schuldiner, M., and M. Bohnert. 2017. A different kind of love – lipid droplet contact sites. Biochim Biophys Acta Mol Cell Biol Lipids. 1862:1188–1196.

37. Schulte, U., F. den Brave, A. Haupt, A. Gupta, J. Song, C.S. Muller, J. Engelke, S. Mishra, C. Martensson, L. Ellenrieder, C. Priesnitz, S.P. Straub, K.N. Doan, B. Kulawiak, W. Bildl, H. Rampelt, N. Wiedemann, N. Pfanner, B. Fakler, and T. Becker. 2023. Mitochondrial complexome reveals quality-control pathways of protein import. Nature. 614:153–159.

38. Seifert, G.J. 2018. Fascinating Fasciclins: A Surprisingly Widespread Family of Proteins that Mediate Interactions between the Cell Exterior and the Cell Surface. Int J Mol Sci. 19.

39. Shai, N., E. Yifrach, C.W.T. van Roermund, N. Cohen, C. Bibi, I.J. L, L. Cavellini, J. Meurisse, R. Schuster, L. Zada, M.C. Mari, F.M. Reggiori, A.L. Hughes, M. Escobar-Henriques, M.M. Cohen, H.R. Waterham, R.J.A. Wanders, M. Schuldiner, and E. Zalckvar. 2018. Systematic mapping of contact sites reveals tethers and a function for the peroxisome-mitochondria contact. Nat Commun. 9:1761.

40. Shevchenko, A., H. Tomas, J. Havlis, J.V. Olsen, and M. Mann. 2006. In-gel digestion for mass spectrometric characterization of proteins and proteomes. Nat Protoc. 1:2856–2860.

41. Sickmann, A., J. Reinders, Y. Wagner, C. Joppich, R. Zahedi, H.E. Meyer, B. Schonfisch, I. Perschil, A. Chacinska, B. Guiard, P. Rehling, N. Pfanner, and C. Meisinger. 2003. The proteome of Saccharomyces cerevisiae mitochondria. Proc Natl Acad Sci U S A. 100:13207–13212.

42. Sorger, D., K. Athenstaedt, C. Hrastnik, and G. Daum. 2004. A yeast strain lacking lipid particles bears a defect in ergosterol formation. J Biol Chem. 279:31190–31196.

43. Sun, L.L., M. Li, F. Suo, X.M. Liu, E.Z. Shen, B. Yang, M.Q. Dong, W.Z. He, and L.L. Du. 2013. Global analysis of fission yeast mating genes reveals new autophagy factors. PLoS Genet. 9:e1003715.

44. van Zutphen, T., V. Todde, R. de Boer, M. Kreim, H.F. Hofbauer, H. Wolinski, M. Veenhuis, I.J. van der Klei, and S.D. Kohlwein. 2014. Lipid droplet autophagy in the yeast Saccharomyces cerevisiae. Mol Biol Cell. 25:290–301.

45. Vogtle, F.N., J.M. Burkhart, H. Gonczarowska-Jorge, C. Kucukkose, A.A. Taskin, D. Kopczynski, R. Ahrends, D. Mossmann, A. Sickmann, R.P. Zahedi, and C. Meisinger. 2017. Landscape of submitochondrial protein distribution. Nat Commun. 8:290.

46. Wang, Q., Y. Sun, T.Y. Li, and J. Auwerx. 2026. Mitophagy in the pathogenesis and management of disease. Cell Res. 36:11–37.

47. Watanabe, Y., Y. Iwasaki, K. Sasaki, C. Motono, K. Imai, and K. Suzuki. 2023. Atg15 is a vacuolar phospholipase that disintegrates organelle membranes. Cell Rep. 42:113567.

48. Weill, U., I. Yofe, E. Sass, B. Stynen, D. Davidi, J. Natarajan, R. Ben-Menachem, Z. Avihou, O. Goldman, N. Harpaz, S. Chuartzman, K. Kniazev, B. Knoblach, J. Laborenz, F. Boos, J. Kowarzyk, S. Ben-Dor, E. Zalckvar, J.M. Herrmann, R.A. Rachubinski, O. Pines, D. Rapaport, S.W. Michnick, E.D. Levy, and M. Schuldiner. 2018. Genome-wide SWAp-Tag yeast libraries for proteome exploration. Nat Methods. 15:617–622.

49. Yofe, I., U. Weill, M. Meurer, S. Chuartzman, E. Zalckvar, O. Goldman, S. Ben-Dor, C. Schutze, N. Wiedemann, M. Knop, A. Khmelinskii, and M. Schuldiner. 2016. One library to make them all: streamlining the creation of yeast libraries via a SWAp-Tag strategy. Nat Methods. 13:371–378.

50. Zielinska, D.F., F. Gnad, K. Schropp, J.R. Wisniewski, and M. Mann. 2012. Mapping N-glycosylation sites across seven evolutionarily distant species reveals a divergent substrate proteome despite a common core machinery. Mol Cell. 46:542–548.

51. Zinser, E., and G. Daum. 1995. Isolation and biochemical characterization of organelles from the yeast, Saccharomyces cerevisiae. Yeast. 11:493–536.

